# Plant Bioengineering Atlas: A Knowledge Graph of Genes, DNA Constructs, and Plant Traits

**DOI:** 10.64898/2026.08.21.746270

**Authors:** Khan A. Yawar, Van C. Nguyen, Akmaljon Makhmudov, Stanton Martin, David J. Weston, Lianhong Gu, Gerald A. Tuskan, Xiaohan Yang

## Abstract

Plant bioengineering has generated tens of thousands of genotype-to-phenotype relationships, but this knowledge remains fragmented across narrative literature and difficult to use computationally. Inconsistent descriptions of DNA constructs, host species, and traits, including variable species names, omitted regulatory elements, and inconsistent gene symbols, impede data reuse, comparative analysis, and design-build-test-learn cycles. Here, we present the Plant Bioengineering Atlas, a literature-mined, ontology-grounded knowledge graph assembled from the PubMed Central (PMC) Open Access Subset using an artificial intelligence (AI)-aided extraction pipeline. A large language model parsed open-access primary research articles to generate structured, automatically extracted, evidence-linked records of engineered genes, modification types, promoter-gene-terminator constructs, host species, target traits, and reported phenotypes, with every record traceable to its source. The current release (v1.1) contains 14,358 evidence-linked records encompassing 6,998 distinct lead gene symbols across 436 plant species from 6,452 papers published between 2000 and 2026. Analysis of the open-access literature shows that experimental efforts are concentrated in a limited set of model and crop species, with disease and pathogen resistance representing the most commonly engineered trait class and constitutive regulation, especially through the CaMV 35S promoter, being the predominant gene expression control strategy. We organize these data into a knowledge graph linking genes, constructs, species, and traits; provide access through an interactive web portal and a versioned repository; and propose an AI-compatible reporting framework for community evaluation. The Plant Bioengineering Atlas provides a foundation for data-driven retrieval, retrospective comparison and hypothesis generation in AI-aided plant biodesign.

## 1. Introduction

Plant genetic engineering has transformed plant science and crop improvement by enabling targeted tests of gene function, expression control, and trait modification. Since the first transgenic plants were produced using *Agrobacterium tumefaciens* as a natural vector ^1, 2^, the field has accumulated a vast experimental record of how individual genes, whether introduced or edited, reshape plant phenotypes ^3^. The advent of programmable nucleases, especially CRISPR-Cas systems, accelerated this trajectory by making targeted modification routine across model and crop species ^4–7^. In parallel, the discipline has shifted toward biosystems design, or biodesign, in which organisms are engineered through iterative design-build-test-learn (DBTL) cycles rather than through one-trait-at-a-time interventions ^3, 8, 9^. Realizing this vision at whole-plant scale requires not only improved tools for writing DNA, but also a comprehensive, computable record of existing engineered constructs and their resulting phenotypes. That record, however, is distributed across thousands of primary research articles. A typical paper reports the engineered gene, the regulatory parts used to express it, the host species, and the resulting phenotype in free text, figures, and supplementary tables, using descriptions that vary widely in completeness and terminology. Foundational results, such as the constitutive activity of the cauliflower mosaic virus (CaMV) 35S promoter ^10^ and the reconstruction of the provitamin A pathway in rice endosperm to address vitamin A deficiency ^11^, have long been regarded as canonical. These results are straightforward in their interpretation but difficult to aggregate, compare, or query at scale. The result is a rich but fragmented knowledge landscape: the same gene may be engineered dozens of times, in different species, and driven by different promoters, yet these instances remain difficult to assemble into a coherent, machine-readable resource. Such a resource is needed because the most useful questions span many studies, for example, which genes have been engineered for the same trait in more than one species, which regulatory parts recur across constructs, or where improving one trait has come at the cost of another. At present, Answering these questions requires tracing evidence across primary research articles.

This fragmentation is the type of obstacle that data standards and structured knowledge resources are designed to address. The FAIR guiding principles establish that research outputs should be Findable, Accessible, Interoperable, and Reusable by machines as well as humans ^12^, while biological ontologies support interoperability by providing controlled vocabularies and logical relationships among entities. The Gene Ontology established this approach, in which entities are annotated with a shared vocabulary whose terms carry defined relationships to one another, for gene function ^13, 14^, and the Planteome project extends it to plants through the Plant Ontology (PO) and the Plant Trait Ontology (TO), enabling species-neutral annotation of plant structures, developmental stages, and traits ^15, 16^. What remains missing is a resource that applies these standards to the central objects of plant bioengineering (i.e., engineered genes, DNA constructs, and resulting traits) and links them with explicit provenance.

Two developments now make such a resource feasible. First, the growth of open-access publishing has placed a substantial fraction of the primary literature within reach of automated processing ^17, 18^. Second, large language models (LLMs) have shown promise for extracting structured information from scientific text at scale ^19, 20^. Together, these developments make it possible to transform the narrative record of plant engineering into the structured relationships required for biodesign.

Here we introduce the Plant Bioengineering Atlas (PBA), a knowledge graph that combines AI-aided literature extraction with ontology-based standardization to create a computable map of plant genetic engineering. We describe the screening and extraction pipeline; characterize a corpus of 14,358 records linking genes, constructs, and traits across 436 species; organize these relationships as a knowledge graph; and make them accessible through an interactive web portal; and test, in a time-split analysis, whether the graph anticipates relationships reported later. We use the corpus to identify patterns within this open-access corpus, including the concentration of studies in a few model species, the pervasive use of the CaMV 35S promoter, and frequent absence of regulatory parts from the extracted construct descriptions. Finally, we propose an AI-compatible reporting framework that would make future plant bioengineering reports machine-readable at publication, supporting reuse of reported results for AI-aided biodesign.

## 2. Methods

The Plant Bioengineering Atlas was assembled in two phases (Supplementary Figure S1). We first mined the open-access literature to create a structured table of plant genetic-engineering experiments and then transformed that table into an ontology-grounded, browsable knowledge graph. All steps after literature retrieval were scripted and, apart from LLM inference, deterministic. LLM output was constrained to a fixed schema, and each extracted record was checked against the source text; subsequent annotations were generated with fixed rule sets and controlled vocabularies rather than additional generative inference.

### 2.1. Data curation and preprocessing

Source articles were obtained from the PubMed Central (PMC) Open Access Subset, distributed as a public Amazon S3 bucket ^21^. Each article version was stored under a prefix named for its PMC accession and version number and included an XML file encoded in the Journal Article Tag Suite (JATS) format and a derived plain-text rendering. At the time of access (29 May 2026), the subset contained 8,969,757 article versions (8,833,423 accessions). We retrieved the plain-text representations by anonymous batch transfer and excluded the deprecated legacy partition. Because the full subset exceeded the storage and file-count limits of the computing cluster, the corpus was processed in shards: each batch was copied to node-local scratch space, screened, and deleted, while only articles that passed screening and their source XML were retained on persistent storage.

Articles reporting plant bioengineering were identified with a purpose-built classifier implemented entirely in the Python standard library. The classifier applied a deterministic cascade of seven rules in a fixed order (minimum length of 500 words; article-type label; review-style language; section structure; plant vocabulary; non-plant dominance; bioengineering vocabulary) and stopped at the first failed rule, thereby assigning each excluded article a single, interpretable rejection reason. The rules are grouped below in four filters: (1) Primary-research filter. Articles with header or body labels, or narrative cues, characteristic of reviews, mini-reviews, perspectives, opinions, commentaries, editorials, case reports or protocols were excluded. (2) Structure filter. At least two canonical IMRaD (Introduction, Methods, Results and Discussion) section headings were required, excluding narratives without the structure of an experimental study. (3) Plant-subject filter. The body had to contain at least two distinct terms from a curated plant vocabulary spanning generic botanical terms, higher taxonomic ranks, and model and crop species in common and binomial forms. A parallel non-plant vocabulary was scored against the same text, and articles dominated by non-plant terms were excluded to suppress incidental botanical mentions in animal, microbial and clinical studies. (4) Bioengineering filter. At least two distinct terms were required from a bioengineering vocabulary covering genome editing, transgenesis and plant transformation, synthetic biology and metabolic engineering, regulatory cassettes and selectable markers, and plant cell and tissue culture. At all stages, matching used word-boundary-anchored regular expressions and counted distinct surface forms rather than raw occurrences, preventing a single high-frequency term from dominating a decision.

Default thresholds (at least two plant-vocabulary hits, two bioengineering-vocabulary hits, two primary-research headings, and a minimum article length of 500 words) were set as a broad first filter because only a small fraction of PMC articles report plant bioengineering; articles with unusual terminology, missing section headings, short formats or few vocabulary matches can be missed. The length threshold was set to filter out stub entries, abstract only deposits, and malformed files, which are common in the subset and too short to support reliable extraction; it was not intended to exclude genuinely short research articles. Applied to the full corpus, the classifier retained 68,300 article versions (67,774 accessions), approximately 0.76% of the input. Excluded versions failed first on article type (4,582,619), plant vocabulary (2,090,822), section structure (1,141,594), bioengineering vocabulary (354,191), length (327,837), non-plant dominance (213,375) or review-style language (191,019) (Supplementary Note 1). For the 490 accessions with more than one retained version, only the lowest-numbered version was processed. Among the articles that contributed records, the JATS metadata link 10 to a correction and 4 to an expression of concern, and none to a retraction; a check against Crossref and Retraction Watch (27 October 2026) likewise found no retracted or withdrawn article. All post-publication notices are listed in the released article table, and no DOI is shared by two contributing articles. Each retained article was saved in its original text and XML formats, named by accession and version, and accompanied by a per-article report recording the decision, the determining rule, and the underlying term counts. These reports provide an auditable trail of every screening decision. Two experts evaluated the screening rules and the LLM gate described in Section 2.2 on random samples of 300 and 200 decisions (Supplementary Table S1.2). The screening rules correctly rejected 208 of 210 sampled rejected articles; with stratum weights, their precision was 0.27 (95% CI 0.18–0.36) and their recall 0.44 (0.21–1.00), as expected for a broad first filter, and the wide recall interval reflects the two missed articles. The gate was more selective: 90 of 100 sampled gate-passed articles were primary plant-engineering studies, giving a precision of 0.90 (0.84–0.95), a recall of 0.72 (0.60–0.87) and an F1 score of 0.80 (0.72–0.89) (Supplementary Table S1.2).

Before extraction, each retained article was reduced to its substantive body. Anchored regular expressions identified section boundaries. Text before the first recognized primary section (Methods, Results, Results and Discussion, Discussion, or Conclusions) and text from the reference list onward, including acknowledgments, author contributions, competing-interest, funding, data-availability, and supplementary sections, were removed. Thus, introductions and other front matter, as well as references and other back matter, were excluded from the model context. Bodies longer than 80,000 characters were shortened to their first and last 40,000 characters to limit the per-article inference cost; this affected 94 of the 8,241 articles that passed the gate described below (1.1%).

### 2.2. AI-aided extraction of structured data from research articles

Structured records were extracted with an open-weight large language model, Qwen3-32B with 4-bit activation-aware weight quantization (AWQ; Qwen/Qwen3-32B-AWQ, revision 0499c3a) ^22^, served locally via vLLM (v0.11.0) ^23^ through an OpenAI-compatible interface. Inference was run on two NVIDIA A40 GPUs using tensor parallelism and greedy decoding (temperature 0, seed 0). These settings were chosen to improve repeatability, but identical output is not guaranteed across runs, software versions, or hardware configurations; the full configuration is given in Supplementary Note 2. Running the model on local hardware kept article text within the host environment and substantially reduced the marginal cost of processing the full corpus.

Each of the preprocessed accessions first went through a keyword prefilter that skipped articles lacking plant or genetic-engineering terms; the remaining articles were evaluated with two sequential model calls. First, a lightweight gate call, which read the first 6,000 characters of the body, determined whether the text was a primary research article that reported plant bioengineering. Its output was constrained to a two-field JSON object containing a Boolean decision and a brief justification. Articles that did not pass the gate were logged and skipped before full extraction. Articles that passed were submitted to an extraction call that returned one structured row for each DNA construct described in the article. In both calls, vLLM guided decoding constrained output at the token level to a JSON schema, forcing each response to follow a fixed template during generation. At each decoding step, only tokens consistent with the schema were permitted, so completed outputs are constrained to the schema. Guided decoding cannot, however, prevent an output from being cut off: 763 articles that passed the gate returned output that could not be parsed (468 reached the output-length limit and 295 had other malformed output) and yielded no records. This approach is more restrictive than simply instructing the model to return JSON, which still permits omitted fields, misnamed keys, incorrect value types, and unexpected fields. The schema fixed field names and value types, restricted the engineering-type field to an enumerated set, required each PMC identifier to match its canonical pattern, and disallowed additional fields.

Each record comprised nine published fields: (1) the host plant species as a Latin binomial (genus-level and hybrid names were kept as reported and never assigned to an assumed species), (2) the engineered gene or genes, each annotated with a symbol, brief functional description, and organism of origin, (3) modification type, such as overexpression, knockout, knockdown, CRISPR-based editing, or RNA interference, (4) the engineering type, distinguishing single-gene from multigene constructs, (5) the DNA construct, (6) a concise phenotype statement, including quantitative values when reported, (7) the proposed molecular mechanism, (8) the DOI, and (9) the PMC identifier of the source article. DNA constructs were encoded with a compact grammar that was both human- and machine-readable. A transcription unit was represented as promoter-gene-terminator, with any unreported element recorded explicitly rather than inferred. Independent transcription units within a multigene construct were joined by triple hyphens, translational fusions by a double colon, and multiplexed CRISPR guide-RNA arrays by a plus sign. Missing information was recorded as “NA”.

To reduce unsupported model output, the schema included a tenth field containing a verbatim evidence quote, which was used internally but omitted from the published table in the Plant Bioengineering Atlas. The requested quote provided a text span against which the record could be checked; matching alone did not establish semantic correctness. The prompt asked that the quote contain the gene name and either the plant species or the specific phenotype or mechanism asserted in the row. Before matching, each quote and the article text given to the model were normalized by standardizing dashes, quotation marks, spaces and ellipses, collapsing repeated spaces and converting to lower case. Quotes shorter than 20 characters were rejected. A quote was accepted as a continuous match when the whole quote occurred in the article text. A quote of at least 60 characters without a continuous match was split into three consecutive segments and accepted as a fallback match when at least two segments, each of at least 15 characters, occurred anywhere in the text. Because the fallback does not require the segments to appear in order or close together, fallback matches are reported separately. Rows whose evidence could not be located were excluded from the table and recorded in a separate audit log. In the archived logs, 16,330 candidate rows gave 10,882 continuous matches, 3,476 fallback matches and 1,972 rejections (Supplementary Note 2); the match type is released for every record. The quote was retained only in that log and was not written to the published table. Together with constrained decoding, this check removes rows that fail the specified text-matching criteria and reduces unsupported output; it confirms that the supporting text exists in the article, not that every field was interpreted correctly. Two plant scientists scored a stratified random sample of 401 released records field by field against the full articles and their supplementary files (Supplementary Tables S1.1, S1.3 and S1.4). The share of records scored correct or partly correct by each expert was 93–100% for host species, gene symbols, gene origin, engineering type, gene part, phenotype, mechanism, DOI and PMCID, 90–92% for modification type, 86–88% for the promoter, and 83–87% for quantitative values; and the corresponding aggregate percentages were similar for continuous and fallback matches (92–94% of field scores). The two experts agreed on whether a field was acceptable or incorrect for 90–99% of records; Cohen’s kappa for the three-category scores was 0.29–0.59 (1.00 for DOI). These values reflect both the concentration of scores in the correct category and disagreements between correct and partly correct (Supplementary Table S1.1). The evaluation also showed that the model had often written the NOS terminator when the article named none. Following the evaluation, terminator names were checked against the article text: a NOS, OCS or 35S terminator was kept only when the article named it and was otherwise recorded as “NA” (3,397 records; Supplementary Note 3).

Extraction was run as a checkpointed batch job on a high-performance computing (HPC) cluster. Completed accessions were recorded continuously, allowing interrupted runs to resume without reprocessing. A total of 67,774 preprocessed accessions entered the screening and extraction workflow. Of these, 19,461 were excluded by the keyword prefilter, and 48,313 underwent LLM-based screening, of which 8,241 passed the gate. Of the gate-passed articles, 1,789 yielded no retained records, including the 763 with unreadable output. Following screening and subsequent quality-control and evidence-verification procedures, the final dataset comprised 14,358 records contributed by 6,452 articles, a mean of 2.2 records per article (Supplementary Figure S1). Every record carries a PMC identifier; 5 records from 5 articles have no DOI because their JATS XML contains none, and 23 extracted DOIs that differed from the JATS value were corrected to it. The resulting nine-column table served as the primary dataset for all subsequent analyses.

### 2.3. Development of the web portal

The extracted nine-column table was transformed into a self-contained web application, in which every analytical field was precomputed. The browser therefore performs only retrieval, filtering, and layout, without a server, database, or network dependency at runtime. The transformation comprised a sequence of deterministic annotation steps implemented in the Python standard library. Each step produced a record-aligned cache that was merged with the source table during the final build. Cache alignment was verified before assembly to guard against row-order drift. Free-text phenotypes and mechanisms were mapped to agronomic trait classes with curated term lexicons, and each class was anchored to a canonical Trait Ontology (TO) term ^15^. Plant structures and tissues were also grounded in the Plant Ontology (PO) ^15, 16^. At least one TO-grounded agronomic trait class was assigned to 7,287 records (50.8% of the dataset); 2,507 of them carry a directly matched TO term and 4,780 an inferred class-level TO term (ontology releases: TO 2026-01-14, PO 2026-01-09, PECO 2025-12-02, PATO 2025-05-14).

To enable cross-species comparisons, engineered genes were assigned to comparative-genomics groups. Gene symbols and identifiers were resolved to PLAZA 5.0 orthogroups ^24^ using the PLAZA identifier-conversion and gene-family (ORTHOFAM) tables for dicot and monocot species. This procedure assigned an orthogroup to 4,557 records (31.7%); the remaining records involve a host species not covered by PLAZA 5.0 (2,283), a gene symbol or identifier that could not be resolved (6,643), or no lead gene or only a reporter or marker gene (875) (Supplementary Table S4). Because genes were looked up in the host genome, the orthogroup of a transgene from another species is that of a putative host homolog. Transcription factors were assigned to PlantTFDB families ^25^ through two complementary routes: a locus-level join between PLAZA and PlantTFDB gene identifiers, and a curated mapping from gene symbols to families. The two routes agreed for 592 of 610 jointly labeled records (97.0%); the locus-based assignment was treated as authoritative when they disagreed. This precedence rule also helps resolve a common ambiguity in plant transcription-factor nomenclature. Many transcription-factor families are named after the cis-regulatory DNA elements they bind, so the protein and its target DNA motif have nearly identical names. Examples include DREB (DRE), CBF (C-repeat), ERF (ethylene-responsive element), ABF/AREB (ABRE), TGA (TGACG), GATA, HSF (heat-shock element), and Trihelix/GT (GT element). In addition, CBF can denote either C-repeat binding factor (AP2/ERF family) or CCAAT-binding factor (NF-Y family). To avoid these naming conflicts, gene-locus information was prioritized whenever available; symbol matching was used only for records lacking locus identifiers.

Several additional attributes were derived from the source fields by fixed rules rather than additional generative inference: (1) manipulation type (gain, loss, or edit of function), (2) effect direction (increase, decrease, or mixed), with reporter-only constructs assigned none, (3) application category, spanning environmental security and the bioeconomy, (4) construct components (promoter, terminator, tag, selectable marker and assembly strategy, parsed from the construct string), and (5) quantitative effects reported in the phenotype. Construct completeness was assessed in two steps. First, missing promoter or terminator information was recorded as “NA”, and a construct was considered complete when every transcription unit named both a promoter and a terminator (Supplementary Note 3). Second, reported promoters and terminators were matched to the promoter (10 classes) and terminator (4 classes) categories listed in Table 1. Constructs were considered resolvable when both elements matched these categories. Reported elements that did not fit the predefined categories were counted as reported but unresolved, making the resolvable estimate a conservative measure of regulatory-part resolvability in the extracted records. The earliest publication date found in each article’s JATS XML was used to determine the publication year. Each article was assigned to the country of its first author’s first listed affiliation, or to that of the article’s first affiliation when the first author had none. The country was read from the country field of the affiliation in the JATS XML or, when this was absent, from the last country name in the affiliation text, and name variants were merged (Supplementary Note 5). The source table and all record-aligned annotations were embedded, together with the client-side application code in a single HTML file. The interface provides complementary modules: a faceted browser backed by an in-memory inverted index for full-text search; an applications view; a catalog of DNA constructs, each linked to its source article by DOI or PMCID; trait-centered and gene-centered views; a trade-off finder; and the knowledge graph described below.

**Table 1.** Promoters and terminators reported in DNA constructs (number of records whose construct string matches each class)

| **Name** | **Description** | **Number of records** |
|---|---|---|
| **Promoter** |  |  |
| CaMV35S | Cauliflower mosaic virus 35S; strong constitutive promoter, widely used in dicots | 5,268 |
| Ubiquitin (Ubi-1 / UBQ) | Maize or Arabidopsis ubiquitin promoter; strong constitutive promoter, commonly used in monocots | 688 |
| U6 / U3 (Pol III) | RNA polymerase III promoters (e.g., AtU6, OsU3) used to transcribe CRISPR guide RNAs | 385 |
| Native / endogenous | The target gene's own (native) promoter | 360 |
| Inducible (Dex/XVE /heat-shock) | Chemically or thermally inducible systems (dexamethasone-GVG, beta-estradiol-XVE, or heat-shock) | 129 |
| Seed-specific | Seed- or endosperm-preferential promoters (napin, glutelin, oleosin, phaseolin) | 67 |
| RbcS / Cab | Light-dependent promoters active in photosynthetic tissues (Rubisco small subunit; chlorophyll a/b-binding) | 61 |
| Actin (Act1) | Plant actin promoter; constitutive, common in monocots | 24 |
| rd29A | Abiotic-stress-inducible promoter (responsive to drought, cold, and ABA) | 14 |
| Root-specific | Literal 'root-specific' class; an expanded search including named root-preferential promoters identified 23 records (Supplementary Table S7) | 1 |
| **Terminator** |  |  |
| NOS terminator | Agrobacterium nopaline synthase polyadenylation signal; the most widely used terminator | 1,074 |
| 35S terminator | Cauliflower mosaic virus 35S terminator | 124 |
| OCS terminator | Agrobacterium octopine synthase terminator | 66 |
| rbcS terminator | Rubisco small-subunit gene terminator | 1 |

### 2.4. Construction of the knowledge graph

The knowledge graph represents the dataset as a network of entities that recur across plant biodesign studies. It consists of six views, each linking two kinds of entity that occur together in the released records. Nodes are lead genes, gene families, transcription-factor (TF) families, agronomic trait classes and host species, each identified by its name; genes that share a symbol in different species therefore share a node. Two nodes are linked by an undirected edge when at least one record contains both, and the edge weight is the number of such records. For multigene records, only the lead gene enters the gene views, while the full gene list remains in the record.

The network can be examined at several levels of abstraction: gene-to-trait, gene-family-to-trait, TF-family-to-trait, species-to-trait, gene-family-to-species and gene-to-species projections. For comparative analyses, users can select two to five traits; the application partitions the contributing genes into shared and trait-specific sets and displays them as an UpSet plot, an intersection matrix, and a trait co-occurrence network. Selecting an intersection reveals the underlying genes, their associated TF families and reported promoters, and their common gene families; orthogroups contain related genes across species and do not by themselves indicate a shared pathway or conserved function.

The same record-level associations support trade-off screening. A gene is flagged as a defense–growth candidate when its records include both a biotic-defense trait class (disease or pest) and a growth or yield class, and as an abiotic–growth candidate when an abiotic-stress class occurs together with a growth or yield class. These flags mark genes whose source articles merit inspection; they are not demonstrated trade-offs. For each gene, the portal also shows a literature-support level based on the number of source articles, the number of host species and the share of records that report the most common direction of effect (Strong: at least 4 articles, 2 host species and 70% agreement; Moderate: at least 2 articles and 60% agreement; otherwise Limited). These thresholds are conventions and do not assess study quality.

Node coordinates were computed with a deterministic force-directed layout that was allowed to settle once and then fixed, rather than animated continuously. The rendered graph supports interactive focus, neighbor expansion, zooming, panning, and export of both the current view and its underlying records.

### 2.5. Time-split test of graph-based hypothesis generation

To assess whether rankings based on earlier records identify relationships reported in later records, we rebuilt it from records published up to a cutoff year and asked how many of its highly ranked, not-yet-reported links were first reported afterwards. The analysis plan, scoring rules and code (Supplementary Note 7) were fixed before ranking and evaluation, after an initial count of later-reported links used to select the tasks and identifier level, and the rankings were locked with SHA-256 checksums before evaluation. Two tasks were defined over the 18 agronomic trait classes, with the article as the unit of evidence. In the transfer task, a candidate was a combination of gene family, trait class and host species in which the family had been reported for the trait in at least one other host and the host had at least 10 articles, but the family had not yet been reported for that trait in that host. In the gene–trait task, a candidate was a PLAZA 5.0 orthogroup and a trait class not yet reported for it. A candidate was counted as positive if an article published after the cutoff reported it.

Candidates were ranked in five ways: by a simple graph rule specified in advance (transfer: the number of hosts in which the family had been reported for the trait; gene–trait: the highest conditional co-occurrence of the candidate trait with the traits already reported for the orthogroup); by a logistic regression on all graph features, which include counts of earlier studies; by a research-intensity baseline (the number of articles already published for the gene family or orthogroup and for the host–trait pair or trait); by a logistic regression on the research-intensity features alone; and at random. The two regressions were fitted only on data before the cutoff (features up to five years before the cutoff, labels from the following five years). The primary analysis used a 2022 cutoff and the records published in 2023–2026; the primary endpoint was the difference in average precision between the two regressions, with 95% confidence intervals from 1,000 bootstrap resamples of gene families or orthogroups. In sensitivity analyses, we required at least two confirming articles. Applying the same procedure to the full release produced a ranked list of untested candidates (Supplementary Data 3).

## 3. Results

### 3.1. Landscape of plant bioengineering in the open-access literature

The current release of the Plant Bioengineering Atlas comprises 14,358 evidence-linked records extracted from 6,452 primary research articles published between 2000 and 2026 (Figure 1A). The number of articles contributing records generally increased over the first two decades, from fewer than 10 in the early 2000s to an initial peak of 488 in 2019. Counts then fell sharply to 133 and 109 articles in 2021 and 2022, respectively, before rebounding to 432, 671, and 737 articles in 2023, 2024, and 2025. This dip is not apparent in estimates of the screened corpus: a random sample suggested that screening retained similar numbers of articles for 2021 and 2022 as for adjacent years, but only 2.5% and 2.4% of them yielded records, respectively, compared with 8–12% in other years (Supplementary Table S8). The low extraction yields indicate that the counts for 2021 and 2022 should be interpreted cautiously. The three most recent complete years account for approximately 29% of all dated papers.

**Figure 1.**
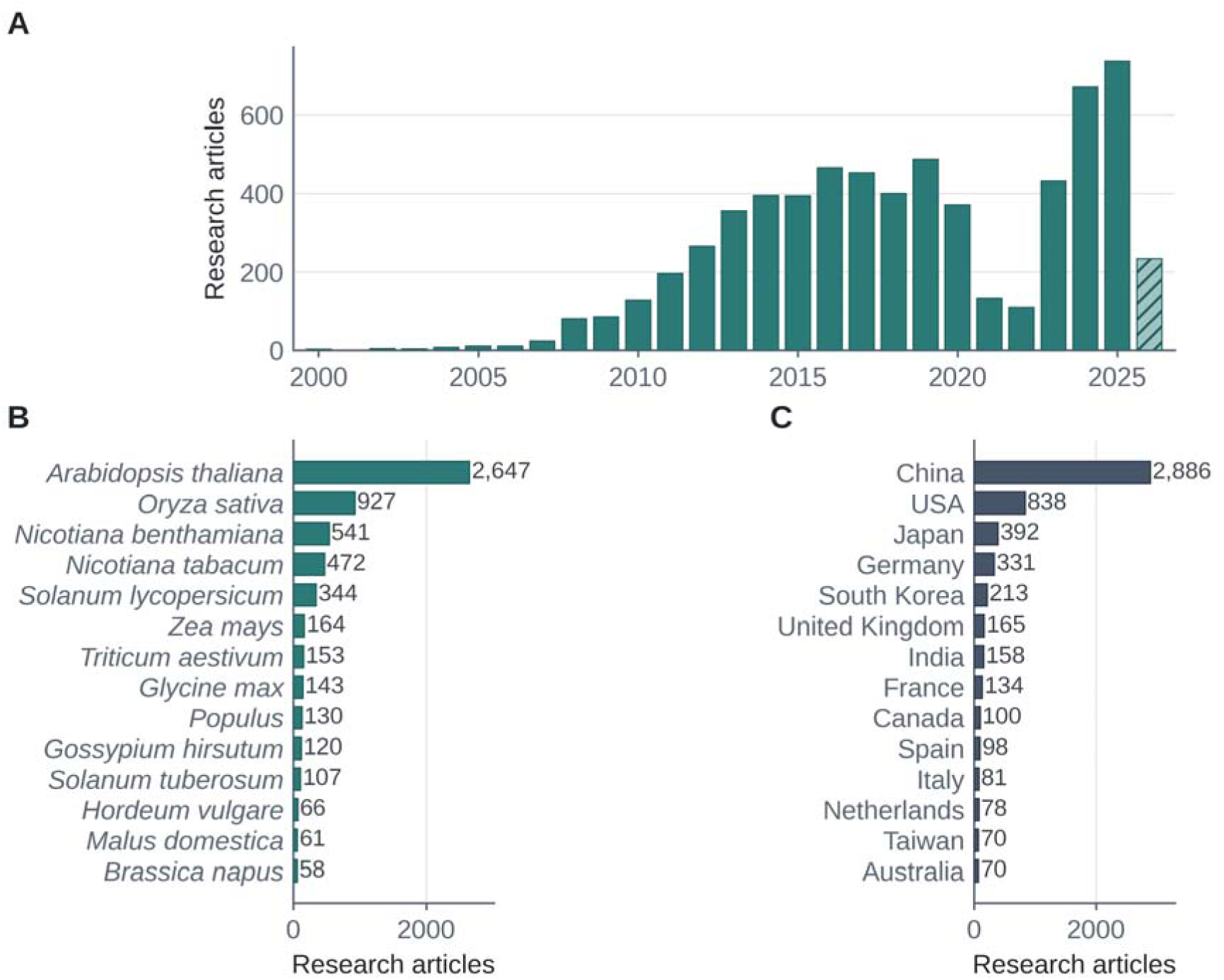
Landscape of plant bioengineering in the PubMed Central open-access literature. (A) Annual number of research articles that contributed records (n = 6,452); 2026 is partial (to 29 May 2026). (B) Number of articles per host label (an article can report several hosts); *Populus* is a genus-level aggregate. (C) Number of articles by country of the first author’s affiliation (6,385 of 6,452 articles resolved).

The 2026 data are partial (234 articles) because the corpus was assembled midway through the year. Within this corpus, plant engineering studies were concentrated in a small number of model species, led by *Arabidopsis thaliana* (2,647 papers; ∼41%), *Oryza sativa* (927; ∼14%), *Nicotiana benthamiana* (541; 8.4%), and *N. tabacum* (472; ∼7%). Major crop and feedstock species followed, including *Solanum lycopersicum* (344 papers; ∼5%), *Zea mays* (164; ∼3%), *Triticum aestivum* (153; ∼2%), *Glycine max* (143; 2.2%), *Populus* spp. (genus aggregate; 130; ∼2%), *Gossypium hirsutum* (120; ∼2%), and *S. tuberosum* (107; ∼2%) (Figure 1B). The geographic distribution of the corpus was similarly concentrated. First-author affiliations mapped 6,385 of 6,452 papers (99.0%) to 67 countries. China (2,886 papers; ∼45%) and the United States (838; ∼13%) accounted for more than half of all resolved affiliations, followed by Japan (392; ∼6%), Germany (331; ∼5%), and South Korea (213; ∼3%) (Figure 1C); 2,796 papers list affiliations in more than one country. This concentration varied by crop: First-author affiliations in China accounted for 52–74% of papers for most major crops, including 65% of rice and 74% of cotton, while potato was a notable exception at 17%, with European groups predominating. Among papers assigned to the United States, 49% concerned Arabidopsis. These first-author affiliation patterns describe differences in host-species representation within this corpus. Because geographic analysis assigns articles by first-author affiliation, we do not attempt to infer underlying national policy, funding, or public-attitude effects, which would require data outside the scope of this resource.

### 3.2. Engineering approaches

Overexpression or ectopic expression was reported in 4,592 of the 6,452 articles (71.2%), followed by heterologous or foreign-gene expression (970; 15.0%), knockout or knockdown (803; 12.4%), gene editing (769; 11.9%), complementation (645; 10.0%) and RNA interference or silencing (596; 9.2%); an article can use several approaches (Figure 2A). At the record level, overexpression accounts for 8,424 records (58.7%) (Supplementary Table S6). The share of papers using genome editing increased over the past decade, from less than 5% before 2015 to 8% in 2017, 13–14% in 2019–2020, and 20–21% in 2023–2025 (Figure 2B), indicating increasing representation of genome-editing approaches within this corpus.

**Figure 2.**
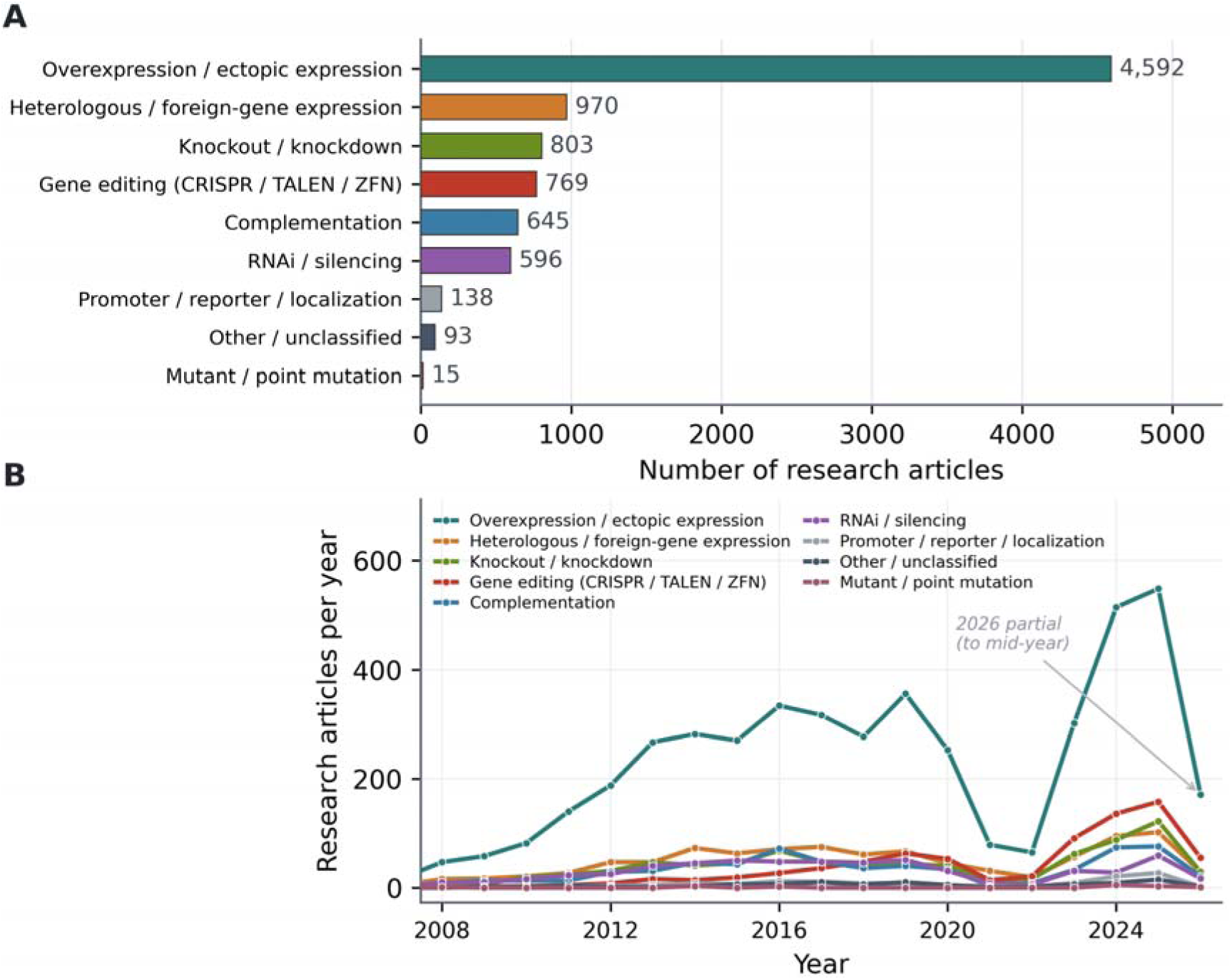
Distribution of plant bioengineering articles in PubMed Central by engineering method. **(A)** Number of articles by engineering-method category (n = 6,452; an article can appear in several categories). **(B)** Annual number of articles by engineering-method category; 2026 is partial.

### 3.3. Genes

The 6,452 papers describe engineering efforts involving 6,998 distinct lead-gene symbols, counted after taking the first gene of each record, ignoring letter case and excluding placeholders and free-text phrases. Because the same symbol can be used in different species, we also paired each symbol with the organism of origin stated in the record, which gives 8,079 symbol–organism pairs; 694 symbols occur with more than one organism of origin, and 2,247 resolve to exactly one PLAZA 5.0 gene model (Supplementary Table S4). Counts of lead-gene symbols per host broadly followed the distribution of articles among hosts: *A. thaliana* accounted for 3,342 engineered genes, more than four times the number in either of th next two model species, *N. benthamiana* (768 genes) and *N. tabacum* (529 genes) (Figure 3A). Other model species were sparsely sampled, with only a few dozen engineered genes each, including *Physcomitrium patens* (61 genes), *Marchantia polymorpha* (39 genes), *Chlamydomonas reinhardtii* (33 genes), and *Brachypodium distachyon* (21 genes). Among crop and feedstock species, rice led with 1,088 engineered genes, followed by tomato (*Solanum lycopersicum*; 398), wheat (198), maize (189), soybean (168), and poplar (157) (Figure 3B). Other crops, such as cotton (135), potato (107), apple (79), and barley (76), were each represented by fewer than 150 genes, indicating lower representation of these hosts within the Atlas. Most genes were engineered in only one or two species; a minority were engineered in more than four host species, including the *Bacillus thuringiensis* insecticidal gene Cry1Ac (10 species), the systemic defense regulator NPR1 (8), the monolignol-pathway enzyme COMT (7), the flavonoid enzyme CHS (6), and the pigmentation and stress-related genes MYB75, PSY1, and WRKY40 (6 each) (Figure 3C). Transcription factors (TFs) were among the most frequently engineered gene classes. Among records assigned to a PlantTFDB family ^25^, the WRKY (268 records), MYB (188), bHLH (161) and ERF (127) families predominated, along with the MIKC-type MADS-box (116), NAC (112) and bZIP (97) families.

**Figure 3.**
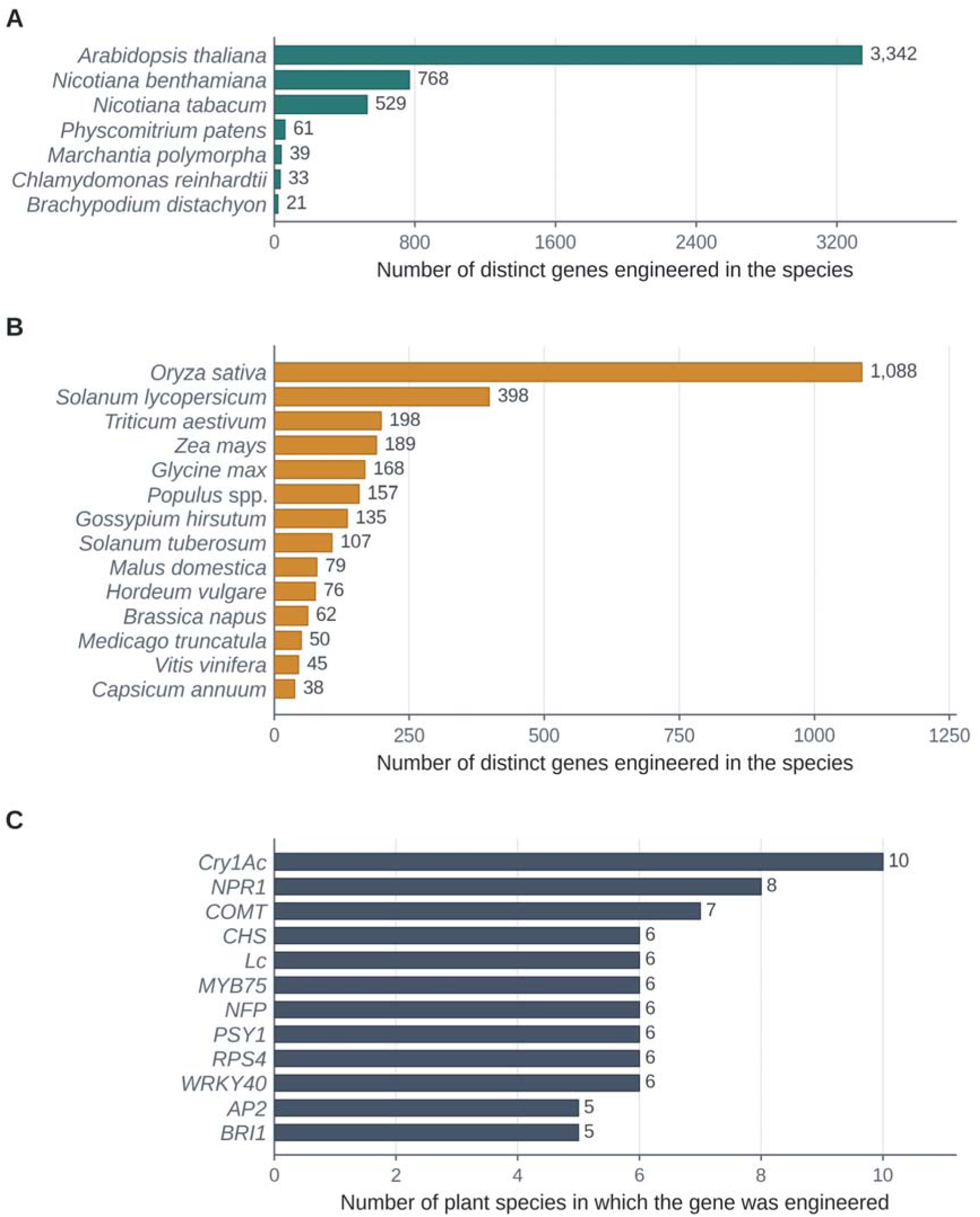
Numbers of genes engineered in representative plant species. (A) Number of distinct lead-gene symbols engineered in model species. (B) Number of distinct lead genes engineered in crop species; Populus spp. is a genus-level aggregate. (C) Genes engineered in the largest number of host species (reporter, marker and tool genes excluded). Each record counts toward its first host species. Cry1Ac: *Bacillus thuringiensis* insecticidal crystal protein Cry1Ac. NPR1: NONEXPRESSOR OF PATHOGENESIS-RELATED GENES 1. COMT: caffeic acid O-methyltransferase. CHS: chalcone synthase. Lc: Leaf color. MYB75: MYB domain transcription factor 75. NFP: Nod Factor Perception. PSY1: phytoene synthase 1. RPS4: RESISTANT TO PSEUDOMONAS SYRINGAE 4. WRKY40: WRKY DNA-binding transcription factor 40. AP2: APETALA2. BRI1: BRASSINOSTEROID INSENSITIVE 1.

### 3.4. DNA constructs

A functional expression construct generally requires a promoter and terminator flanking a coding sequence or other functional component. Construct descriptions were available for 14,185 of 14,358 records (98.8%). Overall, 3,822 records (26.6%) specified a promoter and a terminator for every transcription unit, whereas 10,536 (73.4%) were missing at least one regulatory element (Figure 4A); 855 records of the 14,185 (6.0%) with construct descriptions named a promoter and a terminator that both matched predefined regulatory-part classes. Of the incomplete descriptions, 7,832 specified promoters only, 37 terminators only and 2,494 neither, and 173 records contained no construct description. Because constructs were extracted from the main text only, these values describe the extracted main-text representations. In an automated search of the article XML for 300 records with incomplete constructs, no terminator-related term was found for 270 of the 297 (90.9%) of the records lacking a terminator (Supplementary Table S3). In the expert evaluation, the terminator was not stated in the article for 70–78% of the 401 sampled records (Supplementary Table S1.1). The reported promoter repertoire was relatively narrow: the cauliflower mosaic virus (CaMV) 35S promoter dominated (5,268 records), followed by ubiquitin promoters (688), RNA polymerase III U6/U3 promoters used to express CRISPR guide RNAs (385), and native promoters (360). Tissue-specific and inducible promoters together appeared in fewer than 300 records (Table 1; Supplementary Table S7). Terminators were reported less frequently than promoters. The most frequently reported were the *Agrobacterium* nopaline synthase (NOS) terminator (1,074 records), CaMV 35S terminator (124), and octopine synthase (OCS) terminator (66). Most constructs modified a single gene: 12,982 records (90.4%) described single-gene constructs, whereas 1,372 (9.6%) combined two or more genes in one construct; 4 records have engineering type “NA” (Figure 4B).

**Figure 4.**
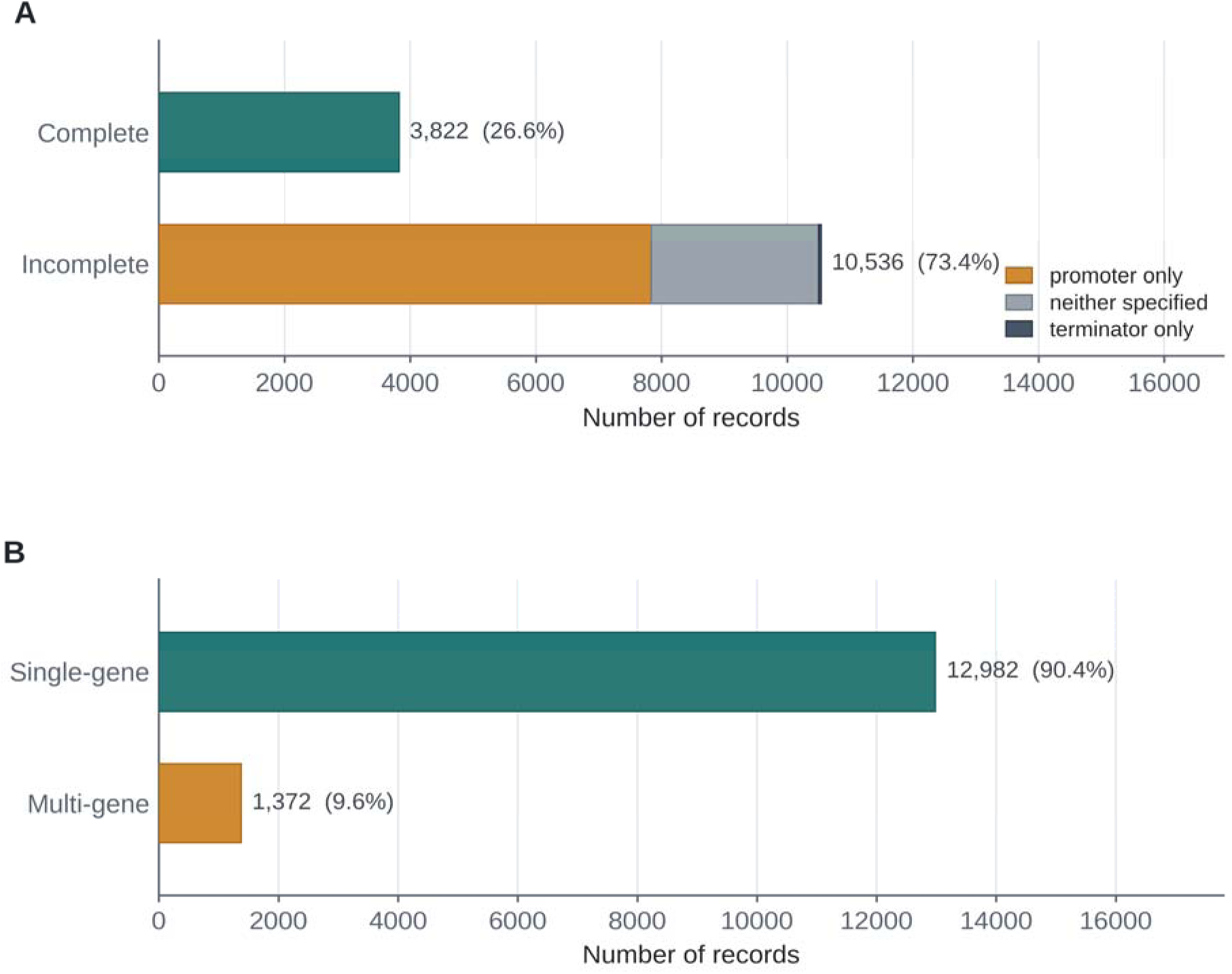
Overview of DNA constructs engineered in plants. (**A**) Records with complete or incomplete promoter-gene-terminator descriptions (n = 14,358; complete = every transcription unit specifies a promoter and a terminator). (**B**) Records describing single-gene and multigene constructs (n = 14,358 overall; 14,354 classified and 4 with engineering type “NA”).

### 3.5. Plant traits

Mapping reported phenotypes with curated lexicons anchored to the Plant Trait Ontology ^16^ assigned at least one of 18 agronomic trait classes to 7,325 records (51.0%) (Figure 5A). Disease and pathogen resistance was the most frequently engineered trait class, followed by flowering and development; yield, grain and biomass; reproduction and fertility; and pigmentation. Individual abiotic-stress tolerance classes were smaller but collectively prominent, including salt (670 records), drought (578), cold (316), heat (202), and herbicide resistance (122). Among trait-annotated records, 5,049 (68.9%) were associated with a single trait class, whereas 2,276 (31.1%) were associated with two or more (Figure 5B), indicating that records often report several traits; this alone does not establish biological pleiotropy. When records were grouped by application, allowing each record to contribute to more than one category, 11,118 records (77.4%) fell into at least one of five plant-biodesign categories: physiological and developmental studies (4,553 records), tool and method development (4,718), environmental and stress resilience (3,568), bioproduct and quality traits (1,674), and industrial or specialty products (310). Within this corpus, physiological studies, methodological tools and stress resilience were more frequently represented than bioproduct and industrial applications; this distribution does not establish the relative research effort across the wider field.

**Figure 5.**
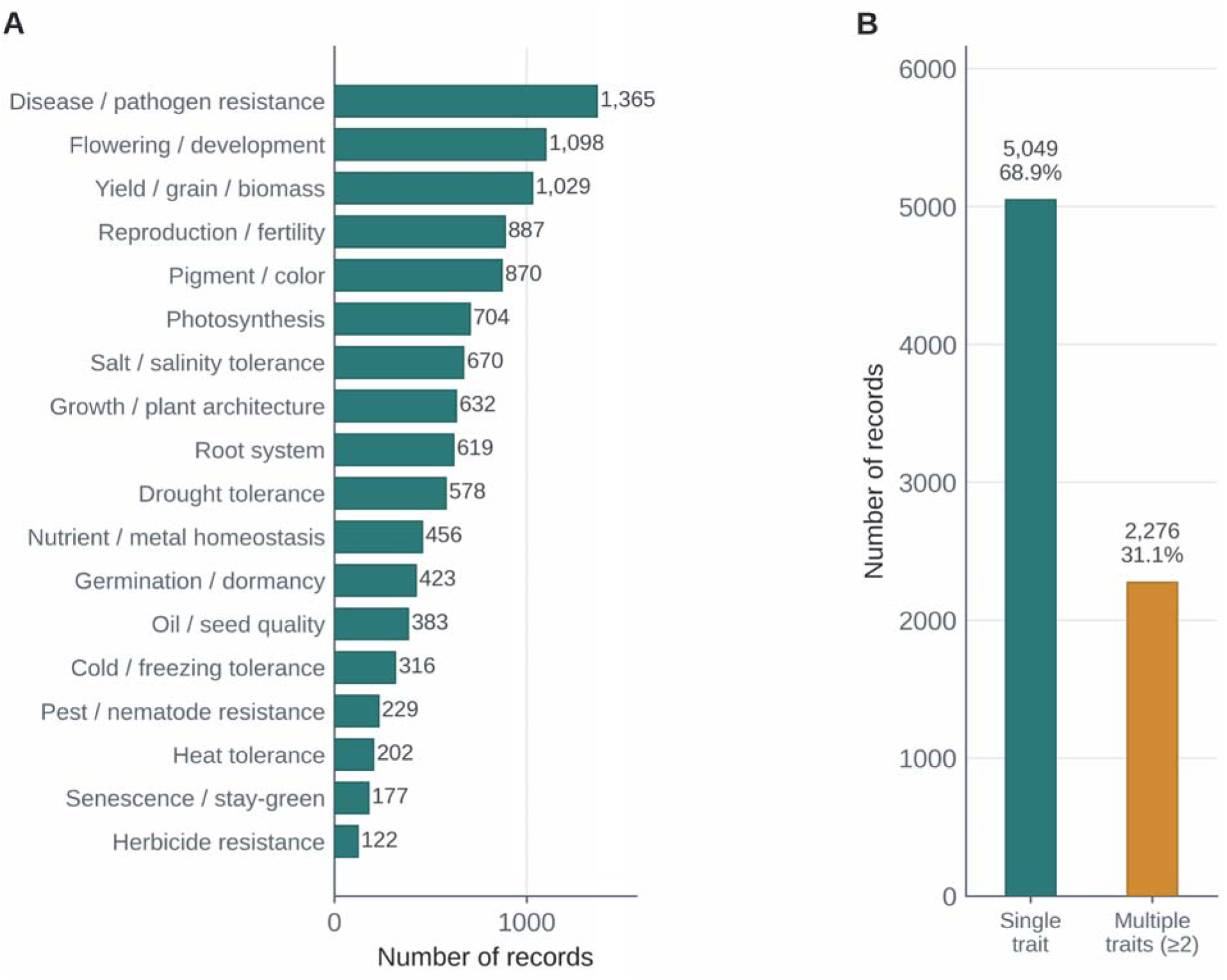
Overview of engineered plant traits. **(A)** Number of records by agronomic trait class (7,325 of 14,358 records carry at least one class; a record can carry several). **(B)** Number of records associated with a single or multiple trait classes.

### 3.6. Trade-offs

Genetic engineering often improves one trait at the expense of another ^3, 8^. Table 2 presents representative, source-linked examples in which a single construct enhances one target trait while compromising another, with the reported outcomes summarized from the cited source articles. These examples span several common categories of engineering trade-off. Growth-defense trade-offs were demonstrated by *BRG8* overexpression, which conferred resistance to rice blast and bacterial blight but induced dwarfism and spontaneous cell death ^26^, and by CRISPR-based knockout of *TaLHP1*, which conferred stripe-rust resistance but caused dwarfism and early flowering ^27^. Stress-development trade-offs were illustrated by *PpCBF1* overexpression, which improved freezing tolerance at the cost of reduced growth and delayed budbreak ^28^; and by *MYB37* overexpression, which enhanced drought tolerance but delayed flowering ^29^. Cross-stress trade-offs were illustrated by silencing *CsGRF04*, which increased drought resistance while reducing salinity and cold tolerance ^30^.

**Table 2.** Examples of engineered DNA constructs that produce trade-offs between plant traits.

| DNA construct<br>(Host species) | Impact on target trait | Unintended impact on another trait | References |
|---|---|---|---|
| CaMV35S-*BRG8Hap*3-EGFP-NES<br>(*Oryza sativa*) | Enhanced resistance to rice blast (*M. oryzae*) and bacterial blight | Dwarfism and spontaneous cell death (growth penalty) | <sup>26</sup> |
| CRISPR/Cas9 knockout of *TaLHP1* (2 gRNAs)<br>(*Triticum aestivum*) | Resistance to stripe rust (*Puccinia striiformis*) | Dwarfism and early flowering | <sup>27</sup> |
| *PpCBF1* overexpression<br>(*Malus domestica*) | Increased freezing tolerance and early dormancy | Reduced stem diameter and height; delayed spring budbreak | <sup>28</sup> |
| CaMV35S-*MYB37*::GFP-NosT<br>(*Arabidopsis thaliana*) | Enhanced drought tolerance (ABA sensitivity, reduced water loss) | Delayed flowering | <sup>29</sup> |
| *CsGRF04* silencing (virus-induced gene silencing, VIGS)<br>(*Citrus sinensis*) | Increased drought resistance | Reduced salinity and cold tolerance | <sup>30</sup> |
| CaMV35S-*OsGL1-3*-NosT<br>(*Oryza sativa*) | Increased tiller number | Reduced grain yield and stunted plant stature | <sup>31</sup> |
| Actin-*vRNA2*(*RGSV p2*)-OCS<br>(*Oryza sativa*) | Increased stem lignin content | Decreased cellulose content and stem mechanical strength | <sup>32</sup> |

### 3.7. Knowledge graph

PBA represents its records as a knowledge graph connecting DNA parts, gene constructs, and plant traits (Figure 6). Engineered genes, their gene families and transcription factor (TF) families, affected plant traits and host species are represented as nodes. An edge connects a gene, or its family, to a trait whenever an extracted record reports that relationship, and edge weight is the number of records (Supplementary Table S6). The graph can be queried at several levels of abstraction. Gene-to-trait, gene-family-to-trait, and TF-family-to-trait projections progressively aggregate individual genes into gene families and TF families, allowing users to examine a trait through either the specific lead-gene symbols associated with it or the broader families to which they were assigned. When two to five traits are selected, the Atlas partitions contributing genes into shared and trait-specific sets and presents them as an UpSet plot, an intersection matrix, and a trait co-occurrence network. This comparative view distinguishes genes engineered for a single trait from genes reported for several traits, as illustrated by the distinct and overlapping gene sets associated with drought and heat tolerance (Figure 7).

**Figure 6.**
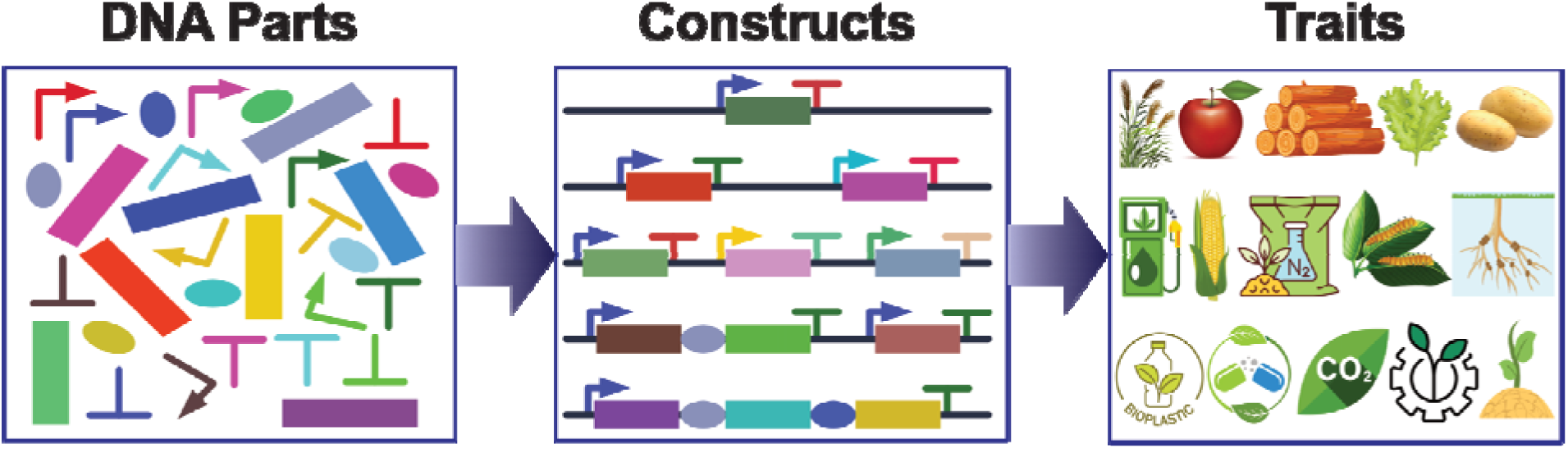
Schematic framework of the plant bioengineering knowledge graph.

**Figure 7.**
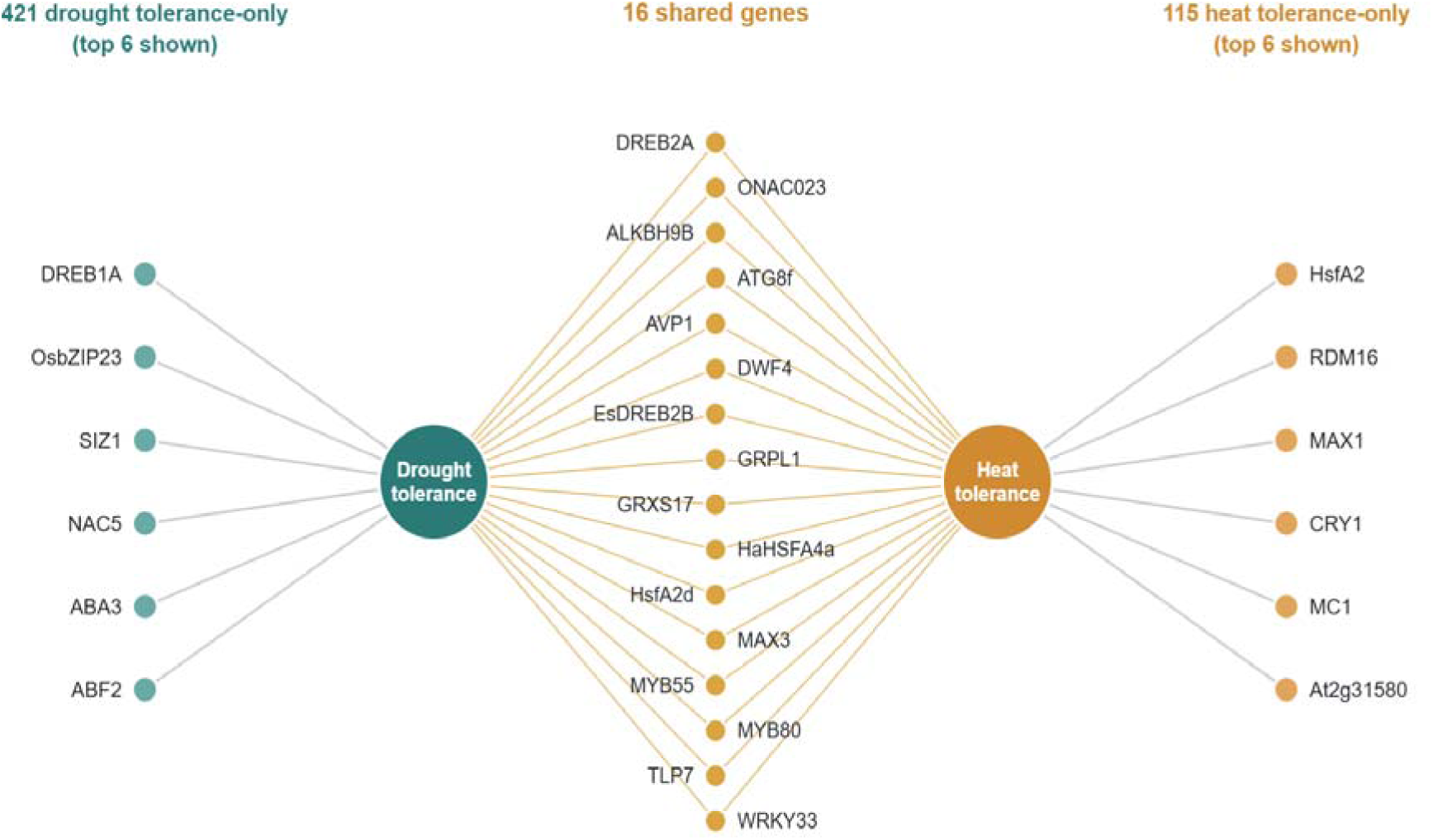
Example knowledge graph for plant bioengineering. The graph shows unique and shared lead genes associated with drought and heat tolerance (gene–trait view; 421 drought-only, 16 shared and 115 heat-only lead genes, of which the six with most records are shown for each trait-only set; reporter and tool genes excluded). DREB1A: dehydration-responsive element-binding protein 1A. OsbZIP23: *Oryza sativa* basic leucine zipper TF 23. SIZ1: SUMO E3 ligase SIZ1. NAC5: NAC-domain transcription factor 5. ABA3: molybdenum-cofactor sulfurase. ABF2: ABRE-binding factor 2. DREB2A: dehydration-responsive element-binding protein 2A. ONAC023: Oryza sativa NAC-domain TF 023. WRKY33: WRKY TF 33. ATG8f: autophagy-related protein 8f. EsDREB2B: *Eremosparton songoricum* DREB2B. HaHSFA4a: *Helianthus annuus* heat-shock TF A4a. MYB55: MYB TF 55. ALKBH9B: AlkB homolog 9B. AVP1: vacuolar H -pyrophosphatase. DWF4: DWARF4. GRPL1: multistress-responsive gene. GRXS17: monothiol glutaredoxin-S17. HsfA2d: heat-shock TF A2d. MAX3: MORE AXILLARY GROWTH 3, carotenoid cleavage dioxygenase 7. MYB80: MYB TF 80. TLP7: thylakoid lipid-associated protein 7. HsfA2: heat-shock TF A2. RDM16: RNA-DIRECTED DNA METHYLATION 16. MAX1: MORE AXILLARY GROWTH 1, CYP711A1. CRY1: cryptochrome. MC1: metacaspase 1. At2g31580: ICARUS1 (ICA1), Thg1-superfamily protein required for cell proliferation at high temperature.

### 3.8. Web interface

To make the Atlas accessible to the broader plant science community, the dataset and analytical views are delivered through an interactive web portal (Figure 8), available at https://fair.ornl.gov/BioDesign/PBA/. The portal is a single self-contained application in which every analytical field is precomputed, so browsing, filtering, and visualization run entirely in the user’s browser without a server, database, or network dependency at runtime. This design keeps the resource fast, portable, and easy to preserve and redistribute as a static archive.

**Figure 8.**
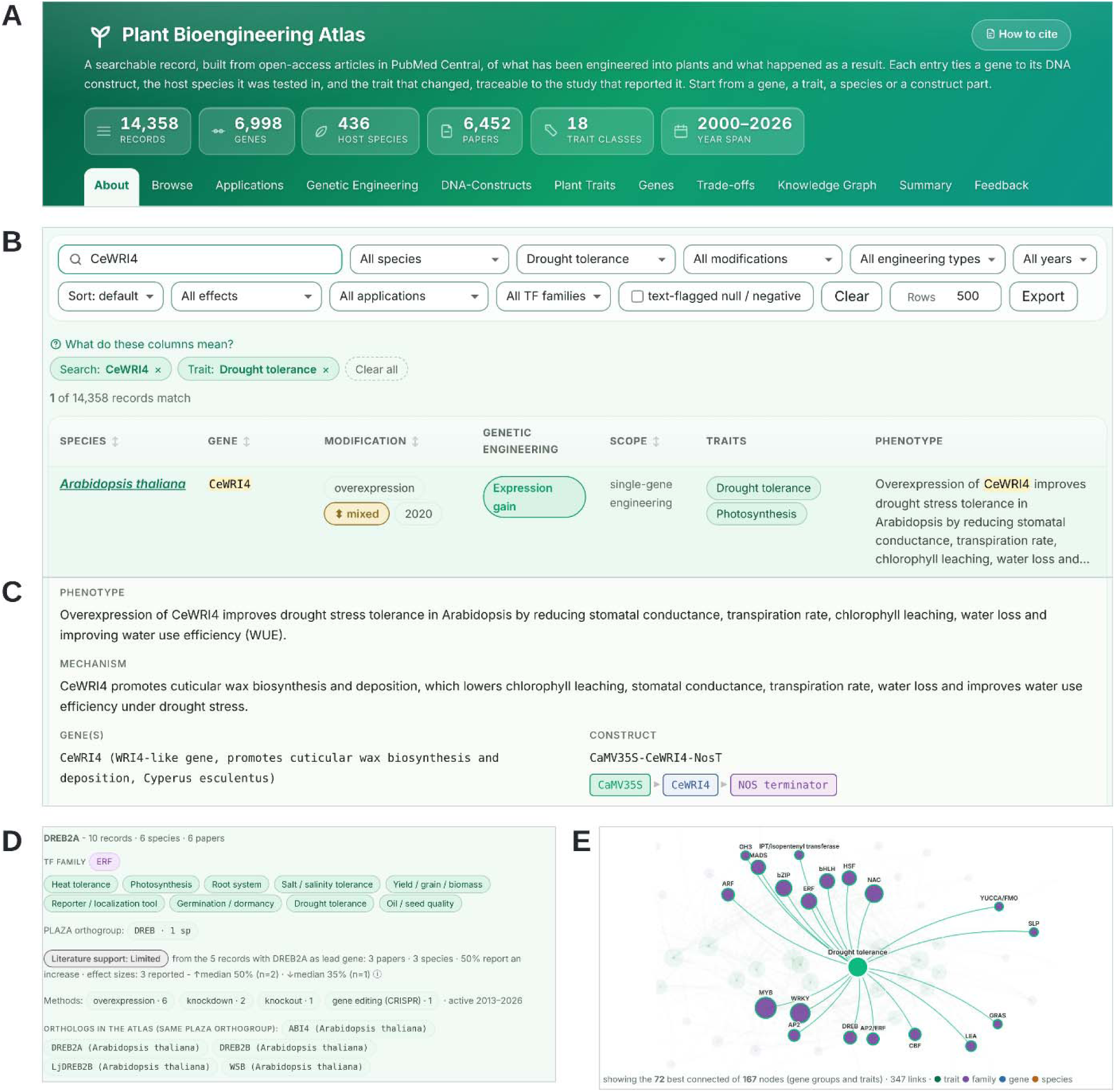
Web portal of the Plant Bioengineering Atlas (data release v1.1). **(A)** Portal header with release statistics and navigation modules. **(B)** Browse view: search and filters for species, trait, modification, engineering type, year, effect, application and TF family, with the matching record. **(C)** The record opened to show the phenotype, mechanism, genes and the parsed DNA construct (promoter, coding sequence and terminator). **(D)** Gene view for DREB2A, with its trait classes, PLAZA orthogroup, literature-support level, methods and orthologs in the Atlas. **(E)** Knowledge graph (gene family–trait view) with drought tolerance selected and its gene-family neighbors highlighted; the 72 best-connected of 167 nodes (gene families and trait classes) are shown. Screenshots of the released portal file, captured and assembled with a browser-automation script.

The interface organizes the data into complementary modules. A faceted browser backed by an in-memory inverted index supports full-text search across all record fields, including genes, species, traits, and constructs, so that a query matches wherever a term appears. An applications view groups records by plant biodesign objective, and a DNA-construct catalog links each construct to its source article by DOI or PMCID. Trait-centered and gene-centered views allow users to enter the corpus from either end of the genotype-to-phenotype relationship; a trade-off finder flags genes reported for both a stress or defense trait and a growth or yield trait; and the knowledge-graph module provides the network and comparative visualizations described above. Every view displays record provenance and allows users to export both the current selection and its underlying data, enabling findings to be traced to the primary literature and reused in downstream analyses.

### 3.9. Anticipating later findings from the graph

Rebuilt from the 9,330 PMC Open Access records published up to the year 2022, the graph yielded 13,793 untested transfer candidates and 17,696 untested gene–trait candidates, of which 162 and 261, respectively, were reported during the period of 2023–2026 (Supplementary Table S9.1). Rankings learned from the graph placed these later reports well above chance. For cross-species transfer, the average precision was 0.138 (95% CI 0.094–0.228) against a base rate of 0.012, and 20 of the 100 top-ranked candidates were reported in PMC Open Access records published between 2023 and 2026 (1.2 expected by chance). For gene–trait links, the average precision was 0.066 (0.047–0.089) against 0.015, and 11 of the top 100 were reported (1.5 expected).

Most of this signal reflects how intensively genes and traits had already been studied: a baseline that used only the number of earlier studies of a gene family, orthogroup, host or trait ranked the later reports similarly (difference in average precision −0.004, 95% CI −0.019 to 0.015, for transfer and 0.003, −0.004 to 0.012, for gene–trait links), and the simple graph rule ranked them less well (Supplementary Tables S9.1 and S9.2). In addition, 216 of the 378 transfers and 240 of the 501 gene–trait links first reported after 2022 involved genes, hosts or trait combinations that were not yet in the graph and could not be ranked. A positive outcome means that a later article reported the link, not that the modification outperformed alternatives. The graph is therefore best used to prioritize candidates with prior support; its top-ranked untested candidates are deposited for prospective testing (Supplementary Data 3).

## 4. Discussion

The Plant Bioengineering Atlas (PBA) provides a large-scale, evidence-anchored, computable account of plant genetic engineering within the PMC Open Access Subset. By coupling AI-aided literature extraction with ontology-based standardization, we transformed the narrative record of thousands of primary research articles into automatically extracted, evidence-linked records describing engineered genes, modification types, DNA constructs, host species, and resulting traits, each traceable to its source publication. Constraining LLM output with token-level guided decoding and requiring a verbatim evidence quote for every record limits plausible claims that are nevertheless unsupported by the source article. These hallucination-suppressing measures yield a structured, auditable corpus. They verify that supporting text exists although they do not necessarily mean that every field is interpreted correctly. Organizing these relationships as a knowledge graph and exposing them through a self-contained web portal turns a fragmented body of literature into a navigable resource for retrieval, retrospective comparison and hypothesis generation in AI-aided plant biodesign. What the Atlas demonstrates so far is the organization of published relationships. In a retrospective time-split test with scoring rules fixed before evaluation (Section 3.9), rankings from the graph anticipated which gene–trait and cross-species links would be reported next far better than chance, although largely because well-studied genes and traits continue to be studied. Showing that the graph can guide new discoveries will require experimental tests of its candidates, which are deposited for this purpose (Supplementary Data 3).

PBA complements, rather than duplicates, existing plant gene-to-trait resources. AgroLD integrates heterogeneous plant genomics, proteomics, and phenomics as a Resource Description Framework (RDF) knowledge graph ^33^; KnetMiner assembles genome-scale knowledge graphs to support evidence-based candidate-gene discovery ^34^; Plant Reactome curates rice pathways and projects them to other species by orthology ^35^; CropGS-Hub assembles genome-wide genotype and phenotype data to support genomic prediction in several major crops ^36^; and literature-derived knowledge graphs predict genes that regulate multiple agronomic traits ^37^. These resources are well suited to questions about gene discovery, pathways, natural allelic variation, and predicted gene–trait relationships. In contrast, PBA documents what has been engineered experimentally, linking genes and traits to the specific DNA constructs used in studies across hundreds of species and thereby informing the design phase of the DBTL cycle.

Methodologically, PBA extends recent work showing that LLMs can recover structured relations from scientific text. Dagdelen et al. ^19^ demonstrated LLM-based extraction of complex records from scientific text, whereas Yao et al. ^20^ evaluated prompt-based extraction of bioregulatory events from rice literature. PBA extends this approach to a broader plant-engineering corpus and adds safeguards tailored to design reuse: guided decoding constrains the output schema, each retained row contains an evidence span located in the article, and downstream annotations are generated with deterministic rules rather than additional generative inference. These decisions favor auditability over maximal recall, a reasonable trade-off when an unsupported promoter, host, or phenotype could propagate into a proposed design. Nevertheless, evidence-span matching verifies textual support, not biological interpretation, and should not be treated as a substitute for expert adjudication. In our two-expert evaluation, most fields were correct or partly correct in more than 90% of records, whereas the promoter, terminator and quantitative values were less reliable (Supplementary Table S1.1). Although developed for plant bioengineering, the pipeline architecture could be adapted to other domains: its components (e.g., screening, extraction anchored to a verbatim quotation, ontology grounding, and delivery as a single self contained file) could be adapted to other fields (e.g., microbial bioengineering, animal bioengineering) with a large open access literature and an established ontology.

PBA has several immediate applications in plant biodesign and biotechnology. Starting from a target trait, researchers can identify repeatedly tested genes, shared regulators, orthologous candidates, and the constructs used to perturb them. Conversely, starting from a gene or regulatory part, users can compare outcomes across host species and engineering strategies. Multitrait graph views can highlight genes reported for both drought and heat tolerance, whereas trade-off screening can identify candidates that may require tissue-specific, inducible, or dosage-tuned expression. Construct catalogs can guide part selection and reveal previously tested combinations, reducing duplicated effort in DBTL cycles. The structured records can also support training and evaluation of information-extraction systems, retrieval-augmented assistants, and predictive genotype-to-phenotype models. In crop improvement, metabolic engineering, and plant-based biomanufacturing, these capabilities could help prioritize experiments with strong prior literature support while drawing attention to underexplored species, traits, and multigene strategies. PBA is intended to support hypothesis generation and design prioritization; final decisions should remain grounded in the relevant experimental context and source publications.

Several limitations arise from the resource’s design. Because extraction was restricted to the PMC Open Access Subset, PBA excludes plant bioengineering research published outside that corpus and inherits the disciplinary, geographic, and temporal biases of openly available literature. Annual article counts therefore do not measure the field’s publication output. The screening rules may also miss relevant articles, and their estimated recall is imprecise (Supplementary Table S1.2). Because one article can contribute several records, record counts are not independent observations; article-level counts are therefore reported alongside record-level counts (Supplementary Table S6). Coverage is also limited by what authors report, so incomplete source descriptions and extraction errors can both produce incomplete construct records, and parts described only in figures, vector maps or supplementary files are not captured. In the graph, genes are identified by symbol alone, so genes that share a symbol in different species are merged into one node, and a transgene from another species is assigned the orthogroup of a putative homolog. The lexicon-based trait and ontology mappings were not validated separately from the record-level expert evaluation. Future releases will seek to broaden coverage through publisher agreements and text- and data-mining licenses for non-open-access literature, extend extraction to full-text PDFs and structured supplementary tables, and refine ontology mappings so that more records can be anchored to canonical trait, structure, and gene-family terms. Periodic re-extraction will keep the resource current as the literature and underlying language models evolve.

Finally, the frequent absence of regulatory parts from extracted construct descriptions motivates changes in how plant bioengineering results are recorded at publication. We therefore propose an AI-compatible documentation framework that makes future reports machine-readable by design. The framework, provided as a JSON Schema with worked examples and a validator (Supplementary Note 6; Supplementary Data 1) and offered for community evaluation rather than as an adopted standard, defines (1) a structured representation of DNA constructs, including promoters, coding sequences, terminators, tags, selectable markers, and assembly logic, expressed through a grammar in which unreported elements are declared explicitly (Figure 9A); (2) a parallel representation of plant traits grounded in the Plant Trait and Plant Ontologies (Figure 9B); and (3) a schema linking each construct to the traits it affects, the host species, and supporting evidence (Figure 9C). Adopted as a lightweight submission supplement, this framework could allow new results to enter resources such as PBA with less reliance on lossy re-extraction, supporting reuse for AI-aided plant biodesign. It would extend machine-readable design representations already developed for synthetic biology, most notably the Synthetic Biology Open Language, which encodes genetic parts and constructs in an ontology-backed, exchangeable form ^38^, and would apply the FAIR principles ^12^ to individual published results. Construct sequence and structure would be referenced as SBOL3 objects rather than encoded again, and plant phenotyping metadata would remain within the scope of MIAPPE (Supplementary Table S5). The proposed profile targets publication-level links among designs, hosts, interventions, outcomes, and evidence, while referring to existing standards for design and phenotyping metadata.

**Figure 9.**
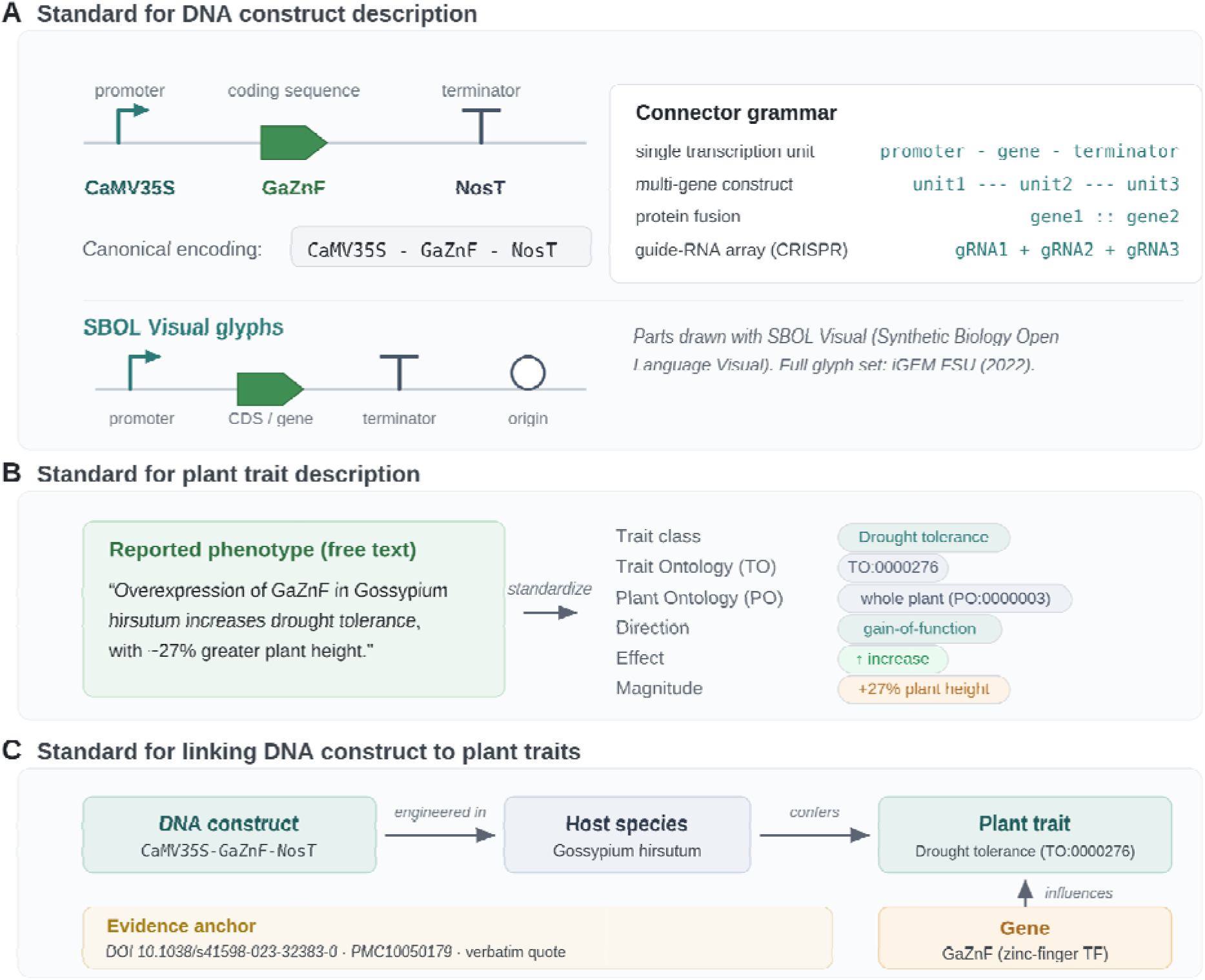
Proposed AI-compatible framework for documenting DNA constructs in plant bioengineering. (**A**) Structured for DNA construct descriptions. (**B**) Structured plant trait descriptions. (**C**) Links between DNA constructs and plant traits.

## Author contributions

X.Y. conceived the idea and revised the manuscript. K.A.Y. performed the data extraction, created the figures and tables, developed the local web portal, and drafted the manuscript. S.M. established the onlin platform for hosting the web portal and revised the manuscript. V.C.N. and A.M. performed the expert evaluation of screening decisions, gate decisions and extracted records. D.J.W., L.G., and G.A.T. reviewed the manuscript and contributed revisions.

## Data availability statement

The Plant Bioengineering Atlas (PBA) is publicly available at https://fair.ornl.gov/BioDesign/PBA/. Data release v1.1 (14,358 records, evidence-check outcome for every record, verbatim evidence quotations for records from CC BY and CC0 articles, article metadata and SHA-256 checksums) is available at https://github.com/khanaliyawar4/Plant_Atlas under CC BY 4.0, and the screening, extraction, annotation, portal and analysis code, prompts, schemas and lexicons are available in the same repository under the MIT license. The repository also holds the reporting-profile files (Supplementary Data 1), the source code (Supplementary Data 2) and the time-split test results (Supplementary Data 3).

## Acknowledgments

This material is based upon work at the Center for Bioenergy Innovation supported by the U.S. Department of Energy (DOE), Office of Science, Biological and Environmental Research (BER) under Contract Number ERKP886. Additional support was provided by the DOE-BER Genomic Science Program through the Plant-Microbe Interfaces (PMI) Scientific Focus Area under FWP ERKP730, and the Genesis Mission Project “A Generative Pre-Trained Transformer for Genomic Photosynthesis” under FWP ERKPA90. Oak Ridge National Laboratory is managed by UT-Battelle, LLC for the U.S. DOE under Contract Number DE-AC05-00OR22725.

## Declaration of generative AI and AI-assisted technologies in the manuscript preparation process

During preparation of this work, the authors used the open-weight large language model Qwen3-32B (Qwen Team) to extract information from research articles, as detailed in the Methods section, and employed generative AI tools to create Figure 9 and improve language clarity. The authors reviewed and edited all outputs as needed and take full responsibility for the content of the published article.

## Supplementary information

Supplementary Notes 1–7, Supplementary Tables S1–S9 and Supplementary Figure S1 are provided in the Supplementary Information file; Supplementary Data 1–3 are deposited in the repository.

