## Supplementary Information for "Plant Bioengineering Atlas: A Knowledge Graph of Genes, DNA Constructs, and Plant Traits"

**Contents**

- Supplementary Note 1: Article screening rules and audit
- Supplementary Note 2: Extraction: preprocessing, model and runtime, prompts, JSON schemas, evidence check
- Supplementary Note 3: DNA-construct grammar, completeness rules and regulatory-part lexicon
- Supplementary Note 4: Annotation layers, knowledge-graph views and portal indicators
- Supplementary Note 5: Country assignment
- Supplementary Note 6: Proposed publication-level reporting profile (schema v0.1.0)
- Supplementary Note 7: Time-split test of graph-based hypothesis generation
- Supplementary Table S1.1: Two-expert evaluation of record fields in 401 sampled records (share of scored records in each category)
- Supplementary Table S1.2: Two-expert evaluation of the screening rules and the LLM gate (stratum-weighted estimates)
- Supplementary Table S1.3: Continuous and fallback matches in the two-expert evaluation (share of field scores that were correct or partly correct)
- Supplementary Table S1.4: Error categories in the two-expert evaluation (one main category per record for each expert)
- Supplementary Table S2: Examples of accepted and rejected candidate rows (random draw, seed 20261026)
- Supplementary Table S3: Automated pre-screen of 300 records with an incomplete construct
- Supplementary Table S4: Annotation coverage by stratum
- Supplementary Table S5: Scope of the proposed reporting profile relative to existing standards
- Supplementary Table S6: Record-level vs article-level shares (paper-level sensitivity)
- Supplementary Table S7: Regulatory-part classes in the released construct strings
- Supplementary Table S8: Screening-retained vs contributing articles by publication year
- Supplementary Table S9.1: Time-split test of graph-based hypothesis generation (cutoff 2022; test records 2023–2026)
- Supplementary Table S9.2: Paired differences between scores in the time-split test (cutoff 2022; test records 2023–2026)
- Supplementary Figure S1: Audited processing flow
- Supplementary Data 1: Reporting-profile schema, examples and validator (reproduced in Supplementary Note 6; deposited in the repository)
- Supplementary Data 2: Source code (the repository)
- Supplementary Data 3: Time-split test results and ranked untested candidates (deposited in the repository)

### Supplementary Note 1: Article screening rules and audit

Input. The PMC Open Access Subset (public S3 bucket pmc-oa-opendata; accessed 29 May 2026) contained 8,969,757 article versions (8,833,423 PMC accessions). Each version's plain-text rendering was classified by a deterministic Python-standard-library classifier (plant_bioeng_filter; full vocabularies and regular expressions in Supplementary Data 2, files rules.py and classifier.py).

Decision rules, applied in this order; the first failed rule is recorded as the rejection reason:

1. Length: fewer than 500 whitespace-delimited words → rejected (stub entries, abstract-only deposits, malformed files).
2. Article type: a non-primary article-type label (review, mini-review, perspective, opinion, commentary, editorial, case report, protocol, etc.) within the first 4,000 characters → rejected.
3. Review language: narrative cues characteristic of reviews/perspectives anywhere in the body → rejected.
4. Structure: fewer than two primary-research section markers (Introduction, Methods, Results, Discussion headings) → rejected.
5. Plant content: fewer than two distinct plant-vocabulary surface forms → rejected.
6. Non-plant dominance: at least three distinct non-plant terms and more non-plant than plant terms → rejected.
7. Bioengineering content: fewer than two distinct bioengineering-vocabulary surface forms → rejected; otherwise retained.

All matching uses word-boundary-anchored, case-insensitive regular expressions and counts distinct surface forms, so a single frequently repeated term cannot satisfy a threshold.

| **Outcome (first failed rule)** | **Article versions** | **% of screened** |
| --- | --- | --- |
| non-primary article type in header | 4,582,619 | 51.1% |
| insufficient plant vocabulary | 2,090,822 | 23.3% |
| missing primary-research sections | 1,141,594 | 12.7% |
| insufficient bioengineering vocabulary | 354,191 | 3.9% |
| text too short | 327,837 | 3.7% |
| non-plant study system dominates | 213,375 | 2.4% |
| review/perspective body language | 191,019 | 2.1% |
| retained | 68,300 | 0.8% |

Versions. The retained 68,300 versions correspond to 67,774 accessions; 490 accessions had more than one retained version. The extraction step processed one version per accession (the lowest version number); 23 of the 6,452 contributing articles had more than one retained version. Corrections and retractions: in the JATS front matter of the 6,452 contributing articles, 10 carry a correction link, 4 an expression-of-concern link and none a retraction link. An external check on 2026-10-27 against Crossref update metadata (6,445 of 6,447 DOIs resolved) and the Retraction Watch database (73,364 entries; matched by DOI and by PMID) found no retracted or withdrawn article; 4 articles carry an expression of concern, 16 a correction, 1 an erratum and 3 an addendum. All notices are listed in the post_publication_notices column of PBA_v1.1_articles.tsv (a14_retraction_check.py). Article types of contributing articles: 6,399 research-article; others: letter 4, case-report 1, brief-report 29, report 1, abstract 12, addendum 1, review-article 3, rapid-communication 1, methods-article 1.

### Supplementary Note 2: Extraction: preprocessing, model and runtime, prompts, JSON schemas, evidence check

#### 2.1 Body preparation and keyword prefilter

Each retained article's plain text was reduced to the span starting at the first line matching a body-start heading (Methods, Results, Results and Discussion, Discussion, Conclusions) and ending at the first back-matter heading (References, Acknowledgments, Author contributions, Conflict/Competing interests, Funding, Supplementary information, Data availability). If no start heading was found, the second half of the article would be used; this fallback was not triggered for any of the 8,241 gate-passed articles. Bodies longer than 80,000 characters were shortened by keeping the first 40,000 and last 40,000 characters (with an explicit truncation marker), so that Methods and the later Results/Discussion were both retained. Of the 8,241 gate-passed articles, 94 (1.1%) exceeded the cap, 70 of which contributed records. The gate call reads the first 6,000 characters of the prepared body.

```
sections:
  body_start_patterns:
    - '^\s*(materials?\s+and\s+methods?|methods?|experimental\s+procedures?)\s*$'
    - '^\s*results?\s*$'
    - '^\s*results?\s+and\s+discussion\s*$'
    - '^\s*discussion\s*$'
    - '^\s*conclusions?\s*$'
  body_end_patterns:
    - '^\s*references?\s*$'
    - '^\s*literature\s+cited\s*$'
    - '^\s*bibliography\s*$'
    - '^\s*acknowledg(e?)ments?\s*$'
    - '^\s*author\s+contributions?\s*$'
    - '^\s*conflict\s+of\s+interest\s*$'
    - '^\s*competing\s+interests?\s*$'
    - '^\s*funding\s*$'
    - '^\s*supplementary\s+(information|materials?|data)\s*$'
    - '^\s*data\s+availability\s*$'
  max_chars: 80000
```

A keyword prefilter then skipped articles whose front-matter article type is a non-research type, or whose full text contains no plant-system term or no genetic-engineering term (configuration below).

```
prefilter:
  skip_article_types:
    - review
    - mini[- ]?review
    - perspective
    - opinion
    - commentary
    - editorial
    - correspondence
    - book\s+review
    - meeting\s+report
    - news
    - erratum
    - corrigendum
    - retraction
  require_plant_terms:
    - plant
    - arabidopsis
    - oryza
    - solanum
    - nicotiana
    - zea\s+mays
    - triticum
    - glycine\s+max
    - medicago
    - populus
    - brachypodium
    - chloroplast
    - leaf
    - stomata
    - phloem
    - xylem
  require_engineering_terms:
    - overexpress
    - knock[- ]?out
    - knock[- ]?down
    - knockdown
    - transgenic
    - crispr
    - cas9
    - cas12
    - guide\s+rna
    - grna
    - sgrna
    - rnai
    - amiRNA
    - hairpin
    - t-dna
    - agrobacterium[- ]mediated
    - transformation
    - gene\s+editing
```

#### 2.2 Model and runtime

| **Item** | **Value (source)** |
| --- | --- |
| Model | Qwen3-32B, 4-bit AWQ; Hugging Face repository Qwen/Qwen3-32B-AWQ, revision 0499c3ac83fdef8810b907a23894ba91e95eddd8 (repository head since 21 May 2025, i.e. the only revision obtainable at download in May 2026) |
| Quantization | AWQ, 4 bits, group size 128, zero point, GEMM layout (repository config.json); loaded by vLLM with the awq_marlin kernel (vLLM log) |
| Inference server | vLLM 0.11.0, OpenAI-compatible API; tensor parallel = 2; dtype float16; max_model_len 32,768; gpu_memory_utilization 0.92; enforce_eager; prefix caching; seed 0; structured-output backend auto (xgrammar); reasoning parser qwen3 (vLLM logs) |
| Software | Python 3.11.11; PyTorch 2.8.0+cu128; transformers >=4.55.2,<5; numpy <2.3 (job logs; environment recipe in Supplementary Data 2) |
| Hardware | 2 x NVIDIA A40 (48 GB); NVIDIA driver 560.35.05; RHEL 8 (ISAAC cluster, University of Tennessee) |
| Decoding | temperature 0.0, top_p 1.0; gate: max 256 tokens, reasoning off; extraction: max 4,096 tokens, reasoning on; JSON-schema-guided decoding for both calls; 8 concurrent articles |

Greedy decoding gives repeatable output on an identical software and hardware stack; it does not guarantee bitwise-identical output across GPU types, kernels, batch composition or library versions, and we make no cross-environment reproducibility claim.

#### 2.3 Prompts (verbatim)

**GATE_SYSTEM**

```
You are a strict triage classifier for biology papers. You return ONLY a
single JSON object. No prose, no markdown fences, no explanation.

Decide whether the article excerpt is BOTH:
  (a) a PRIMARY RESEARCH article — i.e. it presents original experimental
      results. Reviews, mini-reviews, perspectives, opinions, commentaries,
      editorials, news pieces, and protocol-only papers are NOT primary
      research.
  (b) reports PLANT BIOENGINEERING work — i.e. one or more genes are
      overexpressed, knocked out, knocked down, edited (CRISPR/TALEN/ZFN),
      silenced (RNAi/amiRNA), or otherwise modified IN a plant species, and
      the paper measures the resulting effect on the plant.

Return:
  {"qualifies": true,  "reason": "<=200 chars"}     // both (a) and (b) hold
  {"qualifies": false, "reason": "<=200 chars"}     // otherwise

The reason is for human auditing. Keep it short and concrete.

```

**GATE_USER_TEMPLATE**

PMCID: {pmcid}
DOI: {doi}
ARTICLE TYPE (front-matter, if any): {article_type}

--- BODY EXCERPT (first ~6000 chars; may be truncated) ---
{body_excerpt}
--- END EXCERPT ---

Return your JSON now.

**EXTRACT_SYSTEM**

```
You are an expert plant-bioengineering curator. Given the main body of a
primary research article, you extract every DNA construct used to engineer
a plant species in that paper, and for each construct emit one row of a
structured 10-field record.

CRITICAL OUTPUT RULES — read carefully:

  * Return ONLY a JSON object with one top-level field "rows".
  * Return an empty rows list ({"rows": []}) if the paper is NOT plant
    bioengineering, if you find no construct, or if you cannot ground the
    extraction in verbatim text from the body. NEVER invent rows.
  * If a field is genuinely unknown from the text, use the exact string
    "NA" — do NOT guess. (Exception: "evidence_quote" must always be a
    real verbatim quote; if you have no quote, omit the row entirely.)
  * Tabs and newlines inside string values are forbidden; use spaces.
  * The 9 published fields plus the evidence_quote are described below;
    the JSON schema is enforced by the runtime, so emit valid types.

The 10 fields per row, in this order:

1. "plant_species"      — Latin binomial, e.g. "Arabidopsis thaliana".
                          If multiple species are engineered with the SAME
                          construct, list them comma-separated.

2. "genes"              — formatted EXACTLY as:
                          gene1 (name, definition, source), gene2 (name, definition, source), ...
                          where:
                            * name       = the gene symbol/identifier the paper uses
                            * definition = a short functional description from the paper
                            * source     = the organism/source of the gene (e.g. "Arabidopsis
                                            thaliana", "Escherichia coli", "synthetic")
                          If only one gene, still use this template with one entry.

3. "modification_type"  — one of: "overexpression", "knockout", "knockdown",
                          "gene editing (CRISPR)", "RNAi", "amiRNA",
                          "complementation", "expression of foreign gene", or
                          a short phrase that captures it. Multi-gene papers
                          with mixed types: comma-separate (e.g. "overexpression, knockout").

4. "engineering_type"   — EXACTLY one of:
                            "single-gene engineering"  (one gene engineered in this construct/plant)
                            "multi-gene engineering"   (two or more genes engineered)
                            "NA"                       (genuinely undetermined)
                          The runtime rejects any other string.

5. "gene_construct"     — DNA construct using the GRAMMAR below. If the paper
                          does not state promoter/terminator explicitly, use
                          "NA" in that position rather than guessing.

   Grammar for the construct field:
     * Single transcription unit (TU) / single-gene construct:
           promoter-gene-terminator
       Example: "CaMV35S-AtHB17-NosT"

     * Multi-TU / multi-gene construct: connect TUs with three hyphens "---"
           promoter1-gene1-terminator1---promoter2-gene2-terminator2
       Example: "CaMV35S-GeneA-NosT---UBQ10-GeneB-OcsT"

     * Protein fusion: connect fused protein parts with two colons "::"
           promoter-protein1::protein2-terminator
       Example: "CaMV35S-GFP::AtABF3-NosT"

     * CRISPR gRNA array: connect gRNAs with "+"
           promoter-gRNA1+gRNA2+gRNA3-terminator
       Example: "AtU6-gRNA-FT1+gRNA-FT2-polyT"

     * Use the en-dash "-" as the within-TU separator (ASCII hyphen is fine).

6. "phenotype"          — concise sentence describing the impact on the
                          plant, with numbers/units where the paper gives
                          them. Example:
                          "Overexpression of AtHB17 increases leaf biomass
                           by ~15% in Arabidopsis thaliana."

7. "mechanism"          — the molecular mechanism the paper invokes for the
                          phenotype. Example:
                          "AtHB17 enhances photoprotection by accelerating
                           NPQ relaxation, raising effective quantum yield."
                          Use "NA" if the paper proposes no mechanism.

8. "doi"                — DOI as a bare identifier (no URL prefix).
                          Use the DOI shown in the article header.

9. "pmcid"              — the PMC accession, e.g. "PMC12855588" or
                          "PMC12855588.1". The runtime enforces the pattern.

10. "evidence_quote"    — *** ANTI-HALLUCINATION GUARDRAIL ***
                          A VERBATIM contiguous span (~25-150 words) copied
                          EXACTLY from the ARTICLE BODY shown to you below.
                          Character-for-character, including punctuation,
                          numbers, and units. DO NOT paraphrase. DO NOT add
                          ellipsis. DO NOT summarize. DO NOT translate.
                          DO NOT join non-contiguous sentences.

                          The quote MUST contain at least the gene name AND
                          either (a) the plant species name OR (b) the
                          specific phenotype or mechanism described in this
                          row.

                          If you cannot find a single verbatim span that
                          satisfies this — do NOT emit the row. Return an
                          empty rows list instead. Fabricated quotes are
                          detected by a post-hoc substring check and the
                          row will be discarded.

```

**EXTRACT_USER_TEMPLATE**

PMCID: {pmcid}
DOI: {doi}

Extract every plant-bioengineering DNA construct from the article body
below. One JSON row per construct. If the paper engineers the same gene
into multiple species or with multiple constructs, emit a separate row
for each distinct construct.

REMEMBER: each row's `evidence_quote` MUST appear verbatim in the body
text below. Quotes that paraphrase or summarize will be discarded by
the substring verifier.

--- ARTICLE BODY (main text only; intro/refs/acks already stripped) ---
{body}
--- END ARTICLE BODY ---

Return your JSON object now.

#### 2.4 JSON schemas enforced by guided decoding (verbatim)

**Gate call**

```
{
    "type": "object",
    "properties": {
        "qualifies": {"type": "boolean"},
        "reason":    {"type": "string", "maxLength": 200},
    },
    "required": ["qualifies", "reason"],
    "additionalProperties": False,
}
```

**Extraction call (required fields: the nine published columns plus evidence_quote)**

```
{
    "type": "object",
    "properties": {
        "rows": {
            "type": "array",
            "items": {
                "type": "object",
                "properties": {
                    "plant_species":     {"type": "string", "minLength": 1},
                    "genes":             {"type": "string", "minLength": 1},
                    "modification_type": {"type": "string", "minLength": 1},
                    "engineering_type": {
                        "type": "string",
                        "enum": [
                            "single-gene engineering",
                            "multi-gene engineering",
                            "NA",
                        ],
                    },
                    "gene_construct": {"type": "string", "minLength": 1},
                    "phenotype":      {"type": "string", "minLength": 1},
                    "mechanism":      {"type": "string", "minLength": 1},
                    "doi":            {"type": "string", "minLength": 1},
                    "pmcid": {
                        "type": "string",
                        "pattern": r"^PMC[0-9]+(\.[0-9]+)?$",
                    },
                    "evidence_quote": {
                        "type": "string",
                        "minLength": 20,
                        "maxLength": 1200,
                    },
                },
                "required": _MODEL_FIELDS,
                "additionalProperties": False,
            },
        },
    },
    "required": ["rows"],
    "additionalProperties": False,
}
```

Guided decoding constrains the shape of each returned object; it does not prevent truncated generations. 763 gate-passed articles returned output that could not be parsed (468 ended before the JSON object was closed, i.e. reached the token limit; 295 contained other malformed output) and therefore yielded no records.

#### 2.5 Evidence-quotation check (exact algorithm)

8. Normalization of quote and body: a fixed character map folds en/em/minus dashes to '-', curly quotes to straight quotes, non-breaking/thin spaces to spaces and '…' to '...'; runs of whitespace are collapsed to one space; text is lower-cased. No other normalization is applied.
9. Quotes shorter than 20 characters (before normalization) are rejected.
10. Continuous match: the normalized quote occurs as a substring of the normalized body (the same body text the model received).
11. Fallback (only if no continuous match and the normalized quote has at least 60 characters): the quote is cut into three consecutive pieces of floor(n/3), floor(n/3) and the remaining characters (cut on character positions, not words); each piece is trimmed and counts only if it has at least 15 characters and occurs anywhere in the body. The row is accepted if at least two of the three pieces occur. No order, proximity or same-section constraint is applied, which is why fallback-supported records are reported and evaluated separately.
12. Rows failing both tests are removed and written to the audit log with the diagnostic.

| **Evidence-check outcome (archived run logs)** | **Candidate rows** |
| --- | --- |
| accepted: continuous match | 10,882 |
| accepted: 2-of-3 fallback | 3,476 |
| rejected: no match (0–1 of 3 pieces) | 1,947 |
| rejected: no continuous match and quote < 60 characters | 23 |
| rejected: quote < 20 characters | 2 |
| total candidates | 16,330 |

Released records by evidence outcome: continuous matches accounted for 10,882 records (75.8%), while fallback matches accounted for 3,476 records (24.2%), totaling 14,358 records. Median accepted quote length 296 characters. Examples are given in Supplementary Table S2.

### Supplementary Note 3: DNA-construct grammar, completeness rules and regulatory-part lexicon

Grammar requested from the model (ISO/IEC 14977 EBNF; the model's output is not validated against it, see conformance below):

```
construct    = "NA" | unit , { "---" , unit } ;          (* "---" joins independent transcription units *)
unit         = slot , "-" , payload , "-" , slot ;      (* promoter - payload - terminator *)
slot         = "NA" | part ;                            (* "NA" = not stated in the article body *)
payload      = element , { "::" , element }             (* "::" translational fusion *)
             | grna , { "+" , grna } ;                  (* "+" multiplexed guide RNAs *)
element      = part ;  grna = part ;
part         = character , { character } ;              (* author label as written; may itself contain "-" *)
```

Conformance and completeness. Because author labels can contain hyphens, slot boundaries are recovered heuristically. Rule A (used in the submitted Figure 4A): the first and last hyphen-delimited tokens of the whole string are not 'NA' → complete (5,068; promoter only 6,762; terminator only 41; neither 2,314; no construct string 173). Rule A counts 105 one-token and 830 two-token strings as complete although they contain no terminator slot. Rule B (stricter, per transcription unit): every unit has at least three slots and neither its first nor last slot is 'NA' → complete (3,822, 26.6%; promoter only 7,832; terminator only 37; neither 2,494; no construct string 173). The revised Section 3.4 and Figure 4A use Rule B. Both describe the extracted strings, not the source publications.

**Terminator check.** After the expert evaluation, every NOS, OCS or 35S terminator name was checked against the full article text (body, tables and figure legends) and set to 'NA' when the article did not name it; this changed 3,397 records (atlas/data/terminator_corrections.tsv).

Regulatory-part classes (regular expressions applied to the construct string; verbatim from annotate.py):

```
_TERM=[("NOS terminator",r"\bnos[\s-]?t\b|\btnos\b|\bnost\b|nopaline.*?termin|nos termin"),
       ("35S terminator",r"\bt35s\b|35s[\s-]?termin|camv.*?termin"),
       ("OCS terminator",r"\bocs[\s-]?t\b|\bocst\b|octopine.*?termin"),
       ("rbcS terminator",r"rbcs[\s-]?termin"),("HSP terminator",r"hsp[\s-]?termin")]
_PROMO=[("CaMV35S","constitutive",r"\bcamv\s?35s\b|\b2x?\s?35s\b|\bcauliflower mosaic"),
        ("Ubiquitin","constitutive",r"ubiquitin|\bubi\b|\bubq10?\b|zmubq|maizeubq|\bpubi\b|atubq"),
        ("Actin","constitutive",r"\bactin\b|\bact1\b|osactin"),
        ("U6/U3 Pol III","polIII",r"\bu6\b|\bu3\b|atu6|osu6|tau3"),
        ("rd29A stress-inducible","inducible",r"\brd29a?\b"),
        ("RbcS/Cab light","tissue-specific",r"\brbcs\b|\bcab\b"),
        ("seed-specific","tissue-specific",r"seed[\s-]?specific|napin|glutelin|\bgt1\b|oleosin|conglycinin|phaseolin"),
        ("root-specific","tissue-specific",r"root[\s-]?specific"),
        ("inducible (Dex/XVE/HS)","inducible",r"dexamethason|estradiol|\bgvg\b|\bxve\b|heat[\s-]?shock|\bhsp\b|inducible|tetracyclin|\bdex\b|\bg10-90\b"),
        ("native/own","native",r"native|endogenous|own promoter|promoter of|\bpro\b")]
```

### Supplementary Note 4: Annotation layers, knowledge-graph views and portal indicators

Trait classes: 18 agronomic classes (drought, salt, cold, heat, disease, pest, herbicide, yield, flowering, pigment, oil/seed, nutrient, photosynthesis, reproduction, growth/architecture, root, germination, senescence) plus two non-agronomic classes (reporter/localization tool; molecular farming), assigned by curated regular-expression lexicons over phenotype and mechanism text. Each agronomic class is anchored to Plant Trait Ontology (TO) terms. 7,287 records carry at least one TO-grounded agronomic class; 2,507 carry a directly matched TO term and 4,780 an inferred (class-level) TO term. Plant Ontology terms are assigned to 2,429 records. Ontology releases: pato.obo releases/2025-05-14/pato.obo; peco.obo releases/2025-12-02; po.obo releases/2026-01-09; to.obo releases/2026-01-14 (downloaded 8 June 2026).

PLAZA 5.0 orthogroups (dicots and monocots instances): lead gene symbols or explicit gene-model identifiers are looked up in the identifier-conversion table of the HOST species (symbol, full name, alias, MSU ID); the first matching gene model is used and its ORTHOFAM orthogroup assigned. 4,557 records (31.7%) receive an orthogroup. Reasons for no orthogroup: host species absent from PLAZA 5.0 (2,283); host present but symbol/identifier not resolved (6,643); no lead gene or reporter/marker/tool gene (875). Because the lookup uses the host genome, an orthogroup assigned to a heterologous transgene identifies a host homolog; 1,014 orthogroup-bearing records have a lead gene whose stated organism of origin differs from the host.

Transcription-factor families (PlantTFDB species TF lists, 88 species; 58 families): (i) locus-level join of the PLAZA gene model to PlantTFDB; (ii) a curated symbol-to-family lexicon for records without a locus match. Where both routes fire, they agree in 592 of 610 records (97.0%); the locus route takes precedence. Family assignments additionally pass a non-plant/locus-ID guard and the family adjudication table (Supplementary Data 2).

Lead gene symbol: the first gene of each record; distinct labels are counted case-insensitively after removing placeholders and free-text phrases (6,998 labels). Distinct (label, stated organism of origin) pairs: 8,079; labels with more than one stated organism: 694; labels resolved to exactly one PLAZA gene model 2,247, to more than one 54, not resolved 4,697.

Publication year: earliest publication date in the article's JATS XML.

Knowledge-graph views: six bipartite co-occurrence projections computed in the browser from the released records: gene–trait, gene family–trait, TF family–trait, species–trait, gene family–species and gene–species. Nodes are keyed by label within their type (gene nodes by lead-symbol label, without a species namespace; species by canonical host name). An undirected edge links two nodes when at least one record carries both; edge weight = number of records (record- versus article-weighted edges are compared in Supplementary Table S6). Only the lead gene of a record enters gene views; multigene records therefore contribute their lead gene, while the full gene list remains in the record. The 'Records/link' control is a minimum number of supporting records; 'Max nodes' and 'Links/node' only thin the drawing and do not change reported counts. Exports: CSV/Excel tables of the current view and its underlying records, and an image of the view.

Support level shown on gene pages (labeled 'Evidence' in the v1.1 build of 24 August 2026 and 'Literature support' from the October 2026 interface update): computed from the records of that gene: Strong = ≥4 distinct articles, ≥2 host species and ≥70% of records reporting the same effect direction; Moderate = ≥2 articles and ≥60% direction agreement; otherwise Limited. It counts articles, not records, and summarizes literature support within the Atlas; it is unvalidated and does not assess study quality, sample size or risk of bias.

Trade-off finder: genes (reporters excluded) linked to ≥2 agronomic trait classes are listed; a gene is flagged 'defense–growth candidate' when its classes include a biotic-defense class (disease, pest) and a growth/yield class (yield, flowering, photosynthesis, oil/seed, reproduction, growth, germination, senescence), and 'abiotic–growth candidate' when an abiotic class (drought, salt, cold, heat, nutrient, herbicide) co-occurs with a growth/yield class. Flags are co-occurrence candidates for source-level inspection; they do not test whether effects oppose each other in the same experiment.

### Supplementary Note 5: Country assignment

Rule (a8_country.py): the first listed author in the JATS <contrib-group> is taken; the first affiliation linked to that author (xref to <aff>, or an affiliation nested in the contributor element) is used; if the first author has no linked affiliation, the article's first <aff> is used (105 articles). The country is the <country> element when present, otherwise the last country name found in the affiliation string using a curated dictionary of country names and variants (e.g. 'P. R. China', 'People's Republic of China' → China; 'USA', 'United States' → USA; 'Republic of Korea' → South Korea; 'México' → Mexico; Hong Kong and Taiwan reported separately). Articles with only consortium authors or no affiliation remain unresolved.

Result: 6,385 of 6,452 articles resolved (99.0%) to 67 countries; unresolved 67. Top: China 2,886; USA 838; Japan 392; Germany 331; South Korea 213; United Kingdom 165. Sensitivity: corresponding-author country resolved for 3,782 articles and equals the first-author country in 3,651 of 3,782; 2,796 articles list affiliations in more than one country. Agreement with the country file behind the submitted Figure 1C (which covered 6,217 articles and kept name variants separate): 6,107 of 6,131. No manual geocoding service was used. Validation of the string rule (a8b_country_validation.py): on the 655 articles whose first-author affiliation carries a <country> element, the string rule applied to the affiliation text assigned a country to 631 and agreed with the element for all of them; the remainder were unresolved because the affiliation markup is concatenated without separators, so no article was assigned a wrong country.

### Supplementary Note 6: Proposed publication-level reporting profile (schema v0.1.0)

The profile consists of pba_reporting_profile_v0.1.schema.json (JSON Schema 2020-12), a valid and an invalid worked example, and validate_profile.py. All four files are reproduced below and deposited in the repository (folder reporting_profile; Supplementary Data 1). Required core: schema version; publication DOI (PMCID optional); per record: host taxon with resolution rank; each engineered gene with symbol, organism of origin and a stable identifier or an explicit missing value (not_reported / not_applicable / unresolved); intervention class (controlled list); ordered construct parts per transcription unit with roles (promoter … terminator), where an unknown part must be an explicit missing value rather than 'NA' or a blank; at least one outcome with trait label (TO/PO term optional), direction, comparator and evidence location; numeric values require units. Optional: SBOL3 URI of the design, vector, cultivar/line, conditions, mechanism, evidence text span. The validator reports every violation with its JSON path (the invalid example yields four: NA used for a part, missing gene identifier, value without unit, invalid direction). The profile references SBOL3 for sequence-level design instead of re-encoding it; it is a proposal for community evaluation, not an adopted standard.

**pba_reporting_profile_v0.1.schema.json (verbatim)**

```
{
  "$schema": "https://json-schema.org/draft/2020-12/schema",
  "$id": "pba_reporting_profile_v0.1.schema.json",
  "title": "PBA publication-level reporting profile (proposal, v0.1.0)",
  "description": "Links one engineered design to its host, intervention, measured outcome, context and evidence. Proposed for community evaluation; not an adopted standard and not an SBOL replacement. Construct sequence/topology should be referenced as an SBOL3 object where available.",
  "type": "object",
  "required": ["schema_version", "publication", "records"],
  "additionalProperties": false,
  "properties": {
    "schema_version": {"const": "0.1.0"},
    "publication": {
      "type": "object", "required": ["doi"], "additionalProperties": false,
      "properties": {
        "doi": {"type": "string", "pattern": "^10\\.\\d{4,9}/\\S+$"},
        "pmcid": {"type": "string", "pattern": "^PMC\\d+(\\.\\d+)?$"}
      }
    },
    "records": {"type": "array", "minItems": 1, "items": {"$ref": "#/$defs/record"}}
  },
  "$defs": {
    "missing": {"enum": ["not_reported", "not_applicable", "unresolved"]},
    "curie": {"type": "string", "pattern": "^[A-Za-z][A-Za-z0-9_]*:[A-Za-z0-9_.-]+$"},
    "evidence": {
      "type": "object", "required": ["location"], "additionalProperties": false,
      "properties": {
        "location": {"type": "string", "description": "Section, figure, table or supplementary file in the publication, e.g. 'Methods 2.3', 'Fig. 2B', 'Supplementary Table S1'."},
        "text_span": {"type": "string", "maxLength": 1500}
      }
    },
    "taxon": {
      "type": "object", "required": ["name", "rank"], "additionalProperties": false,
      "properties": {
        "name": {"type": "string", "description": "Scientific name as resolved (author label kept in author_label)."},
        "rank": {"enum": ["species", "subspecies", "hybrid", "genus"]},
        "ncbi_taxon": {"type": "string", "pattern": "^NCBITaxon:\\d+$"},
        "cultivar_or_line": {"type": "string"},
        "author_label": {"type": "string"}
      }
    },
    "gene": {
      "type": "object", "required": ["symbol", "origin_taxon", "identifier"], "additionalProperties": false,
      "properties": {
        "symbol": {"type": "string"},
        "origin_taxon": {"$ref": "#/$defs/taxon"},
        "identifier": {"oneOf": [{"$ref": "#/$defs/curie"}, {"$ref": "#/$defs/missing"}], "description": "Stable gene/locus identifier qualified by the organism of origin, e.g. TAIR:AT5G05410, or an explicit missing value."},
        "author_label": {"type": "string"}
      }
    },
    "part": {
      "type": "object", "required": ["role", "name"], "additionalProperties": false,
      "properties": {
        "role": {"enum": ["promoter", "5UTR", "cds", "gRNA", "tag", "linker", "terminator", "marker", "other"]},
        "name": {"oneOf": [{"type": "string", "minLength": 1, "not": {"enum": ["NA", "na", ""]}}, {"$ref": "#/$defs/missing"}]},
        "identifier": {"type": "string", "description": "Registry/GenBank accession or SBOL3 URI of the part, if available."}
      }
    },
    "transcription_unit": {
      "type": "object", "required": ["parts"], "additionalProperties": false,
      "properties": {"parts": {"type": "array", "minItems": 1, "items": {"$ref": "#/$defs/part"}, "description": "Ordered 5'->3'."}}
    },
    "outcome": {
      "type": "object", "required": ["trait", "direction", "comparator", "evidence"], "additionalProperties": false,
      "properties": {
        "trait": {"type": "object", "required": ["label"], "additionalProperties": false,
                  "properties": {"label": {"type": "string"}, "to_term": {"type": "string", "pattern": "^TO:\\d{7}$"}, "po_term": {"type": "string", "pattern": "^PO:\\d{7}$"}}},
        "direction": {"enum": ["increase", "decrease", "no_change", "mixed"]},
        "value": {"type": "number"},
        "unit": {"type": "string"},
        "comparator": {"type": "string", "description": "e.g. 'wild type Col-0', 'empty-vector control'."},
        "conditions": {"type": "string"},
        "evidence": {"$ref": "#/$defs/evidence"}
      },
      "dependentRequired": {"value": ["unit"]}
    },
    "record": {
      "type": "object", "additionalProperties": false,
      "required": ["host", "genes", "intervention", "construct", "outcomes", "evidence"],
      "properties": {
        "host": {"$ref": "#/$defs/taxon"},
        "genes": {"type": "array", "minItems": 1, "items": {"$ref": "#/$defs/gene"}},
        "intervention": {"enum": ["overexpression", "heterologous_expression", "complementation", "rnai_silencing", "knockout_knockdown", "genome_editing", "base_editing", "prime_editing", "mutant_allele", "reporter_or_promoter_analysis", "other"]},
        "construct": {
          "type": "object", "required": ["transcription_units"], "additionalProperties": false,
          "properties": {
            "grammar_string": {"type": "string", "description": "Compact PBA grammar (Supplementary Note 3), derived from transcription_units."},
            "transcription_units": {"type": "array", "minItems": 1, "items": {"$ref": "#/$defs/transcription_unit"}},
            "vector": {"type": "string"},
            "sbol3_uri": {"type": "string", "format": "uri"}
          }
        },
        "outcomes": {"type": "array", "minItems": 1, "items": {"$ref": "#/$defs/outcome"}},
        "mechanism": {"type": "string"},
        "evidence": {"$ref": "#/$defs/evidence"}
      }
    }
  }
}
```

**example_valid.json (verbatim)**

```
{
  "schema_version": "0.1.0",
  "publication": {"doi": "10.1007/s11103-015-0415-2", "pmcid": "PMC4740368"},
  "records": [{
    "host": {"name": "Arabidopsis thaliana", "rank": "species", "ncbi_taxon": "NCBITaxon:3702", "cultivar_or_line": "Col-0"},
    "genes": [{"symbol": "MYB37", "origin_taxon": {"name": "Arabidopsis thaliana", "rank": "species", "ncbi_taxon": "NCBITaxon:3702"}, "identifier": "TAIR:AT5G23000"}],
    "intervention": "overexpression",
    "construct": {
      "grammar_string": "CaMV35S-MYB37::GFP-NosT",
      "transcription_units": [{"parts": [
        {"role": "promoter", "name": "CaMV 35S"},
        {"role": "cds", "name": "MYB37"},
        {"role": "tag", "name": "GFP"},
        {"role": "terminator", "name": "NOS terminator"}]}]
    },
    "outcomes": [
      {"trait": {"label": "drought tolerance", "to_term": "TO:0000276"}, "direction": "increase", "comparator": "wild type Col-0", "evidence": {"location": "Results"}},
      {"trait": {"label": "flowering time"}, "direction": "increase", "comparator": "wild type Col-0", "evidence": {"location": "Results"}}
    ],
    "evidence": {"location": "Methods; Results"}
  }]
}
```

**example_invalid.json (verbatim)**

```
{
  "schema_version": "0.1.0",
  "publication": {"doi": "10.1000/example"},
  "records": [{
    "host": {"name": "Oryza sativa", "rank": "species"},
    "genes": [{"symbol": "GENE1", "origin_taxon": {"name": "Oryza sativa", "rank": "species"}}],
    "intervention": "overexpression",
    "construct": {"transcription_units": [{"parts": [{"role": "promoter", "name": "NA"}, {"role": "cds", "name": "GENE1"}]}]},
    "outcomes": [{"trait": {"label": "grain yield"}, "direction": "up", "value": 12.5, "comparator": "wild type", "evidence": {"location": "Fig. 3"}}],
    "evidence": {"location": "Results"}
  }]
}
```

**validate_profile.py (verbatim)**

import json, sys
from jsonschema import Draft202012Validator
schema = json.load(open(sys.argv[2] if len(sys.argv) > 2 else "pba_reporting_profile_v0.1.schema.json"))
Draft202012Validator.check_schema(schema)
doc = json.load(open(sys.argv[1]))
errs = sorted(Draft202012Validator(schema).iter_errors(doc), key=lambda e: list(e.absolute_path))
for e in errs: print("INVALID", "/".join(map(str, e.absolute_path)) or "<root>", "-", e.message[:160])
print("VALID" if not errs else f"{len(errs)} violation(s)"); sys.exit(1 if errs else 0)

### Supplementary Note 7: Time-split test of graph-based hypothesis generation

**Purpose and freezing.** The test asks whether rankings computed from the graph anticipate links that are first reported later. The analysis plan, the code (pba_prospective.py) and all parameters were fixed before any record published after the cutoff was read. The prediction step discards those records while reading the release file and writes SHA-256 hashes of the code (57c860d0…edba), the release file (26afd4d7…054a) and every ranking; the evaluation step stops if any hash differs. Before freezing, only the numbers of links first reported after 2022 were counted, to choose the tasks and the identifier level; no score was evaluated on later records, and the code was tested on synthetic data. Prediction and evaluation ran as separate jobs on 9 November 2026.

**Outcomes.** The 18 agronomic trait classes are used, with the article as the unit of evidence. Cross-species transfer: a candidate is a gene family F, trait class T and host H such that F was reported for T in at least one host by the cutoff, H has at least 10 articles by the cutoff, and F was not yet reported for T in H. Gene–trait association: a candidate is a PLAZA 5.0 orthogroup O reported by the cutoff and a trait class T not yet reported for O. A candidate is positive if an article published after the cutoff reports it. Gene families are used for transfer because PLAZA orthogroups (31.7% of records) give only 25 transfer events after 2022.

**Scores.** Graph rule (pre-specified): transfer, the number of hosts in which F was reported for T; gene–trait, the largest conditional co-occurrence P(T | T′) over the traits T′ already reported for O. Graph model: a logistic regression on graph features (transfer: hosts with F–T evidence, articles for F–T, F, H–T, H and F in H, and gene families studied for T in H; gene–trait: largest and summed conditional co-occurrence, share of other orthogroups of the same gene family reported for T, articles for O and T, and traits already reported for O). Research-intensity baseline: log articles for F plus log articles for H–T (transfer) or log articles for O plus log articles for T (gene–trait). Research-intensity model: a logistic regression on these counts only. Random: a seeded hash. Counts are log(1 + n) and standardized; the regressions use a ridge penalty of 1 and are fitted on data before the cutoff only (features from records up to five years before the cutoff, labels from links first reported in the following five years).

**Evaluation.** Primary analysis: cutoff 2022 (9,330 records), test records 2023–2026 (5,028). Primary endpoint: the difference in average precision (AUPRC) between the graph model and the research-intensity model, with a 95% CI from 1,000 bootstrap resamples of gene families (transfer) or orthogroups (gene–trait), seed 20261109. Secondary: AUROC, precision and recall among the top k candidates and the other paired differences. Sensitivity: positives supported by at least two articles.

**Ranked untested candidates.** The same procedure applied to the full release ranks every link not yet reported; the 50 highest-ranked transfers and gene–trait links by the graph model, with their supporting PMCIDs, are in Supplementary Data 3 for comparison with later reports.

**Limitations.** A positive means that a later article reported the link, not that the modification worked better than an alternative; negative results are rarely published. Gene families are coarse, orthogroups cover 31.7% of records, keyword-based trait classes and extraction errors carry into both the features and the outcomes, and no experimental test was performed.

### Supplementary Tables

#### Supplementary Table S1.1: Two-expert evaluation of record fields in 401 sampled records (share of scored records in each category)

| **Field** | **Expert 1: n** | **correct %** | **partly correct %** | **incorrect %** | **Expert 2: n** | **correct %** | **partly correct %** | **incorrect %** | **Cohen's kappa** |
| --- | --- | --- | --- | --- | --- | --- | --- | --- | --- |
| Host species | 401 | 95.0 | 2.2 | 2.7 | 401 | 94.8 | 2.2 | 3.0 | 0.59 |
| Gene symbols | 401 | 80.8 | 18.7 | 0.5 | 398 | 73.1 | 23.9 | 3.0 | 0.34 |
| Gene origin organism | 401 | 75.8 | 17.2 | 7.0 | 385 | 76.9 | 17.9 | 5.2 | 0.46 |
| Modification type | 401 | 75.6 | 16.0 | 8.5 | 399 | 80.7 | 9.0 | 10.3 | 0.49 |
| Engineering type | 401 | 86.5 | 9.2 | 4.2 | 399 | 93.7 | 3.0 | 3.3 | 0.33 |
| Promoter | 401 | 77.3 | 11.0 | 11.7 | 362 | 80.9 | 4.7 | 14.4 | 0.41 |
| Gene part | 400 | 73.2 | 24.2 | 2.5 | 394 | 66.8 | 30.7 | 2.5 | 0.52 |
| Terminator, after correction | 401 | 76.1 | 4.0 | 20.0 | 270 | 60.7 | 2.6 | 36.7 | – |
| Phenotype | 399 | 71.7 | 25.1 | 3.3 | 395 | 69.9 | 27.3 | 2.8 | 0.42 |
| Quantitative values | 293 | 21.2 | 62.1 | 16.7 | 122 | 43.4 | 43.4 | 13.1 | 0.29 |
| Mechanism | 361 | 48.2 | 46.3 | 5.5 | 345 | 58.0 | 38.0 | 4.1 | 0.45 |
| DOI | 401 | 99.8 | 0.0 | 0.2 | 401 | 99.8 | 0.0 | 0.2 | 1.00 |
| PMCID | 401 | 100.0 | 0.0 | 0.0 | 401 | 100.0 | 0.0 | 0.0 | – |

**Notes.** Sample (drawn with seed 20261026): 401 records stratified by evidence outcome (241 continuous, 160 fallback, allocated proportionally to host group × period and drawn by simple random sampling within each stratum). A field is scored against the full article and its supplementary files: correct, matches the article; partly correct, the right entity but incomplete or imprecise; incorrect, wrong or not supported by the article. Records scored 'not applicable' are excluded; for the promoter and the terminator, 'not stated in the article' counts as correct when the record gives 'NA' and as incorrect otherwise. Quantitative values and mechanism were mostly scored partly correct because records give a concise statement and not every number or step in the article; 13–17% and 4–6% were scored incorrect. Terminator, after correction: the corrected records (Supplementary Note 3) re-scored against the same expert judgments. Agreement: the two experts agreed on whether a field was acceptable (correct or partly correct) or incorrect for 93–99% of records for most fields (90% for quantitative values). Kappa is modest because most records were scored correct, which lowers kappa even when agreement is high (host species: 96% agreement, kappa 0.59), and because 519 of the 681 disagreements were between correct and partly correct.

#### Supplementary Table S1.2: Two-expert evaluation of the screening rules and the LLM gate (stratum-weighted estimates)

| **Step** | **Sample** | **Judged primary plant-engineering research** | **Precision (95% CI)** | **Recall (95% CI)** | **F1 (95% CI)** |
| --- | --- | --- | --- | --- | --- |
| Screening rules | 90 retained articles; 210 rejected articles (30 per rule) | 24 of 90 retained; 2 of 210 rejected | 0.27 (0.18–0.36) | 0.44 (0.21–1.00) | 0.33 (0.21–0.48) |
| LLM gate | 100 passed articles; 100 rejected articles | 90 of 100 passed; 8 of 100 rejected | 0.90 (0.84–0.95) | 0.72 (0.60–0.87) | 0.80 (0.72–0.89) |

**Notes.** Samples (drawn with seed 20261026): 200 gate decisions (100 passed and 100 rejected, drawn at random) and 300 screening decisions (30 drawn uniformly at random from all versions failing each rejection rule, by reservoir sampling over the full decision table, plus 90 drawn at random from the retained versions). Because strata are sampled at different rates, precision and recall are estimated with stratum weights equal to the number of decisions in each stratum; the 95% confidence intervals are from 2,000 bootstrap resamples within strata. The screening rules are a broad first filter that keeps any article resembling plant genetic work, so their precision is low by design; the LLM gate then selects the engineering studies. Screening recall rests on two articles: both were rejected for too few bioengineering terms, a rule that rejected 354,191 article versions, so two of 30 sampled articles scale to a large estimated number and the interval is wide. The gate is deliberately strict, which keeps its precision high and lowers its recall. After the two experts reviewed the source articles, their gate and screening decisions were identical for all 200 and all 300 articles (kappa 1.00).

#### Supplementary Table S1.3: Continuous and fallback matches in the two-expert evaluation (share of field scores that were correct or partly correct)

| **Evidence-check outcome** | **Records** | **Expert 1: field scores correct or partly correct** | **Expert 2: field scores correct or partly correct** |
| --- | --- | --- | --- |
| Continuous match | 241 | 92.3% | 93.6% |
| Fallback match | 160 | 91.9% | 92.8% |

#### Supplementary Table S1.4: Error categories in the two-expert evaluation (one main category per record for each expert)

| **Main error category of the record** | **Expert 1: records (%)** | **Expert 2: records (%)** |
| --- | --- | --- |
| Cassette misassembly | 109 (27%) | 139 (35%) |
| Omitted fact | 128 (32%) | 9 (2%) |
| Host/origin confusion | 40 (10%) | 34 (8%) |
| Direction or number error | 30 (7%) | 12 (3%) |
| Mechanism over-interpretation | 29 (7%) | 4 (1%) |
| Symbol ambiguity | 13 (3%) | 10 (2%) |
| Identifier error | 0 (0%) | 1 (<1%) |
| Other | 47 (12%) | 154 (38%) |
| None | 5 (1%) | 38 (9%) |

#### Supplementary Table S2: Examples of accepted and rejected candidate rows (random draw, seed 20261026)

| **Outcome** | **Diagnostic** | **PMCID** | **Host** | **Construct** | **Evidence quote** |
| --- | --- | --- | --- | --- | --- |
| accepted: continuous match | strict | PMC11587779 | Arabidopsis thaliana | NA-PbELF4a-NA | Ectopic overexpression of PbELF4a and PbELF4b in Arabidopsis significantly delayed flowering and suppressed the expression of flowering-related genes. |
| accepted: continuous match | strict | PMC13036091 | Oryza sativa | NA-OsGF14c-NLS-GFP-NA | To verify the translocation during M. oryzae infection, we inoculated OsGF14f-cMyc and OsGF14c-cMyc plants and investigated the nucleocytoplasmic distribution of OsGF14f and OsGF14c. Immunoblot analyses revealed that OsGF14f predominantly accumulated in the nucleus during M. oryzae infection, with a slight increase in the cytoplasm (Fig. 6a). In contrast, OsGF14c exhibited a slight cytoplasmic inc |
| accepted: continuous match | strict | PMC11875497 | Zea mays | NA-cry1Ab-NA---NA-cry2Ab-NA---NA-g10evo-epsps-NA | The transgenic maize ZhedaRuifeng8 (ZDRF8) contains cry1Ab, cry2Ab and g10evo-epsps (a selection marker) gene expression cassettes, conferring resistance to lepidopteran pests and tolerance to glyphosate. |
| accepted: continuous match | strict | PMC5322733 | Arabidopsis thaliana | CaMV35S-RFP::BB-NosT | Overexpression of BB under the control of the 35S promoter leads to strongly reduced growth, and, when crossed with a line overexpressing DA1, this growth inhibition was reversed. |
| accepted: 2-of-3 segment fallback | chunked (2/3) | PMC3676348 | Arabidopsis thaliana | NA-PLDδ-NA | Previous studies reported the generation of a PLDδ (PLDδ, AT4G35790)-knockout mutant (PLDδ-KO) and PLDδ-complemented plants (PLDδ-RC) in the background of the Arabidopsis Wassilewskija (WS) ecotype [37], as well as the production of a line, PLDα1-antisense (PLDα1-AS), that was derived from the Arabidopsis ecotype Columbia (Col) and in which PLDα1 activity was attenuated by antisense-mediated suppr |
| accepted: 2-of-3 segment fallback | chunked (2/3) | PMC4706423 | Arabidopsis thaliana | CaMV35S-CFLAP1SRDX-NosT | To elucidate the function of CFLAP1 and its homologous genes, we used chimeric repressor technology [65–67] to further knock down the activities of CFLAP1 and its homologs by overexpressing an ethylene response factor (ERF)-associated amphiphilic repression (EAR) motif-fused CFLAP1 construct, 35S:CFLAP1SRDX, in wild-type Arabidopsis (Fig 5A). Fifty 35S:CFLAP1SRDX independent transgenic lines were |
| accepted: 2-of-3 segment fallback | chunked (2/3) | PMC11101322 | Vitis vinifera | TRV-VvWRKY5 | The VvWRKY5-pRI (VvWRKY5 overexpression) and VvWRKY5-TRV (virus-induced VvWRKY5 suppression) recombinant vectors were constructed and injected into grape fruits by Agrobacterium-mediated genetic transformation before being treated with a wooden stick. The findings revealed that VvWRKY5 overexpression significantly promoted wounding-induced anthocyanin biosynthesis; conversely, VvWRKY5 inhibition r |
| accepted: 2-of-3 segment fallback | chunked (2/3) | PMC10118128 | Manihot esculenta | pCsCMV-AS2/LOB47 | Silencing of MeASLBD47 reduced the lesion area and enabled differential expression of target genes. |
| accepted: 2-of-3 segment fallback | chunked (2/3) | PMC7448358 | Lycopersicon esculentum | CaMV35S-hairpinRNA-LeGPA1-NosT | The RNAi construct was built by inserting a PCR-generated 220-bp segment of LeGPA1 cDNA (segment 340–560 bp) (the RNAi target) into pGM-T plasmid, followed by transformation into E. coli DH5α. The correct positive clone was named pGM-S... Finally, the pCAS was transferred into Agrobacterium GV3101 through freeze-thawing. |
| accepted: 2-of-3 segment fallback | chunked (2/3) | PMC3634990 | Glycine max | FMV-Glyma08g14550.1a-NA | When gene A25 was overexpressed in transgenic soybean roots, the female index decreased to 42% of the control. |
| rejected | no match (1/3 chunks present) | PMC4333230 | Arabidopsis thaliana | Hsp::Cre---loxP-H2B:tdTomato---RPS5Ap::loxP---GCS1 | To rescue the infertility of sperm cells in gcs1/+ mutants, a construct was designed as shown in Fig. 1 (see Supplementary Table S1 for primers used for construction). Three fragments for Hsp::Cre, loxP-H2B:tdTomato, and RPS5Ap::loxP were amplified from a vector (DKv137; pMDC100/Hsp::Cre-RPS5Ap::loxP-GUS-loxP-H2B-tdTomato-NosT)... The fragments RPS5Ap::loxP, Hsp::Cre, and loxP-H2B:tdTomato were cloned into the |
| rejected | no match (0/3 chunks present) | PMC1781449 | Petunia hybrida | CaMV35S-GUS-OcsT | Infiltration was successful also for detached petunia flowers... with strong positive signal for samples ranging from 15 mm-length buds to fully opened flowers... GUS or GFP signal was not observed in any of the respective negative control plants. |
| rejected | no match (1/3 chunks present) | PMC13031928 | Arabidopsis thaliana | AtU6-gRNA-MGO1-NosT---RPS5a-zCas9i-rbcS-E9---nos-BAR-nosT | The binary vector containing CRISPR/Cas9 reagents used to generate mgo1CRISPR was created with the modular cloning (MoClo) toolkit. Annealed oligonucleotides comprising the spacer sequence targeting MGO1 (At5g55300) (5‘-attgATCATCAGTTCCATCTTCTC-3‘, 5‘-aaacGAGAAGATGGAACTGATGAT-3‘) were cloned using the Esp3I restriction enzyme into the previously described Level1_blank_sgRNAv2_Sp_AtU6_(Position 3) |
| rejected | no match (1/3 chunks present) | PMC13143221 | Arabidopsis thaliana | CaMV35S-CITF1-MYC-NosT | We utilized the citf1-1 mutant... introduced a construct expressing CITF1 fused to a 9×MYC epitope tag under the control of the CaMV 35S promoter (35Spro:CITF1-MYC). In parallel, we generated citf1-1 plants expressing FIT fused to GFP under the same promoter (35Spro:FIT-GFP). These transgenic lines were subsequently crossed to obtain a co-expression line (35Spro:CITF1-MYC + 35Spro:FIT-GFP) in the |
| rejected | no match (1/3 chunks present) | PMC11880317 | Arabidopsis thaliana | proMAG3-GFP-MAG3-NosT | proMAG3:GFP-MAG3 was transferred into the destination vector pHGW59 using LR clonase II (Invitrogen). proMAG3:GFP-MAG3 was transformed into the wild-type (Col-0), mag5-3, mag3-5 or proMAG3:GFP-MAG3 mag3-5 plants with Agrobacterium using the floral dip method61 to generate proMAG3:GFP-MAG3 mag3-5. Expression of GFP-MAG3 by the native promoter successfully rescued the mag3 phenotype related to trans |
| rejected | no match (1/3 chunks present) | PMC4846459 | Arabidopsis thaliana | CaMV35S-SbFNSII-2-NosT | The cDNA of the FNSII-2 gene was overexpressed in transgenic Arabidopsis under the control of the cauliflower mosaic virus (CaMV) 35S promoter. [...] When pinocembrin was present in the medium, a new peak corresponding to chrysin was detected in all five FNSII-2–expressing lines, but not in controls. Plants expressing FNSII-2 converted most of the pinocembrin they absorbed to chrysin, accumulating |

#### Supplementary Table S3: Automated pre-screen of 300 records with an incomplete construct

| **Missing element** | **n** | **term in model input (possible extraction miss; upper bound)** | **term only elsewhere in article XML** | **no such term anywhere in article text** |
| --- | --- | --- | --- | --- |
| terminator | 297 | 24 | 3 | 270 |
| promoter | 114 | 74 | 12 | 28 |

**Notes.** Terms searched: terminator names and generic 'terminator'/polyadenylation; promoter names and generic 'promoter', 'driven by', 'under the control of'. A term in the body does not prove it belongs to this construct, so the first column is an upper bound on extraction misses. Of the 270 terminator-missing records with no terminator term in the article text, 232 come from articles with supplementary files and 119 name a vector in the body (the element may be specified only in a vector map, supplement or cited protocol). In the expert evaluation of 401 records (Supplementary Table S1.1), the terminator was not stated in the article for 78% of records (first expert) and not stated or not applicable for 70% (second expert).

#### Supplementary Table S4: Annotation coverage by stratum

| **Stratum** | **n** | **Agronomic trait class** | **Direct TO term** | **PO term** | **Orthogroup** | **TF family** | **No orthogroup: reporter/no gene** | **No orthogroup: host not in PLAZA** | **No orthogroup: not resolved** |
| --- | --- | --- | --- | --- | --- | --- | --- | --- | --- |
| All records: all | 14358 | 7325 (51.0%) | 2507 (17.5%) | 2429 (16.9%) | 4557 (31.7%) | 1940 (13.5%) | 875 (6.1%) | 2283 (15.9%) | 6643 (46.3%) |
| Host group: Arabidopsis thaliana | 6106 | 2845 (46.6%) | 804 (13.2%) | 1328 (21.7%) | 3740 (61.3%) | 1170 (19.2%) | 377 (6.2%) | 1 (0.0%) | 1988 (32.6%) |
| Host group: Other hosts | 4512 | 2531 (56.1%) | 935 (20.7%) | 582 (12.9%) | 80 (1.8%) | 421 (9.3%) | 318 (7.0%) | 1278 (28.3%) | 2836 (62.9%) |
| Host group: Oryza sativa | 1919 | 1214 (63.3%) | 598 (31.2%) | 406 (21.2%) | 714 (37.2%) | 248 (12.9%) | 70 (3.6%) | 0 (0.0%) | 1135 (59.1%) |
| Host group: Nicotiana benthamiana / N. tabacum | 1821 | 735 (40.4%) | 170 (9.3%) | 113 (6.2%) | 23 (1.3%) | 101 (5.5%) | 110 (6.0%) | 1004 (55.1%) | 684 (37.6%) |
| Publication period: 2019-2026 | 7190 | 3774 (52.5%) | 1415 (19.7%) | 1174 (16.3%) | 1945 (27.1%) | 985 (13.7%) | 345 (4.8%) | 1262 (17.6%) | 3638 (50.6%) |
| Publication period: 2011-2018 | 6477 | 3236 (50.0%) | 1020 (15.7%) | 1118 (17.3%) | 2327 (35.9%) | 862 (13.3%) | 453 (7.0%) | 944 (14.6%) | 2753 (42.5%) |
| Publication period: 2000-2010 | 691 | 315 (45.6%) | 72 (10.4%) | 137 (19.8%) | 285 (41.2%) | 93 (13.5%) | 77 (11.1%) | 77 (11.1%) | 252 (36.5%) |
| Modification class: Overexpression / ectopic expression | 8424 | 4491 (53.3%) | 1661 (19.7%) | 1406 (16.7%) | 2651 (31.5%) | 1334 (15.8%) | 522 (6.2%) | 1298 (15.4%) | 3953 (46.9%) |
| Modification class: Heterologous / foreign-gene expression | 1578 | 537 (34.0%) | 94 (6.0%) | 298 (18.9%) | 444 (28.1%) | 151 (9.6%) | 261 (16.5%) | 278 (17.6%) | 595 (37.7%) |
| Modification class: Gene editing (CRISPR / TALEN / ZFN) | 1205 | 587 (48.7%) | 214 (17.8%) | 172 (14.3%) | 322 (26.7%) | 100 (8.3%) | 54 (4.5%) | 151 (12.5%) | 678 (56.3%) |
| Modification class: Knockout / knockdown | 1121 | 687 (61.3%) | 238 (21.2%) | 160 (14.3%) | 278 (24.8%) | 149 (13.3%) | 8 (0.7%) | 258 (23.0%) | 577 (51.5%) |
| Modification class: Complementation | 865 | 441 (51.0%) | 139 (16.1%) | 207 (23.9%) | 571 (66.0%) | 71 (8.2%) | 3 (0.3%) | 10 (1.2%) | 281 (32.5%) |
| Modification class: RNAi / silencing | 778 | 459 (59.0%) | 143 (18.4%) | 127 (16.3%) | 160 (20.6%) | 64 (8.2%) | 13 (1.7%) | 198 (25.4%) | 407 (52.3%) |
| Modification class: Promoter / reporter / localization | 210 | 74 (35.2%) | 8 (3.8%) | 43 (20.5%) | 84 (40.0%) | 48 (22.9%) | 5 (2.4%) | 38 (18.1%) | 83 (39.5%) |
| Modification class: Other / unclassified | 154 | 40 (26.0%) | 9 (5.8%) | 13 (8.4%) | 36 (23.4%) | 22 (14.3%) | 9 (5.8%) | 49 (31.8%) | 60 (39.0%) |
| Modification class: Mutant / point mutation | 23 | 9 (39.1%) | 1 (4.3%) | 3 (13.0%) | 11 (47.8%) | 1 (4.3%) | 0 (0.0%) | 3 (13.0%) | 9 (39.1%) |

#### Supplementary Table S5: Scope of the proposed reporting profile relative to existing standards

| **Aspect** | **SBOL3** | **MIAPPE** | **Planteome (PO/TO)** | **Proposed profile (v0.1.0)** |
| --- | --- | --- | --- | --- |
| Primary scope | Genetic design: parts, sequences, topology | Plant phenotyping experiment metadata | Controlled vocabularies for plant anatomy, development, traits | Publication-level link: design – host – intervention – measured outcome – evidence |
| Construct | Full design model | Not modeled | Not applicable | Ordered parts per transcription unit + optional SBOL3 URI |
| Missing parts | Absent objects | — | — | Explicit not_reported / not_applicable / unresolved |
| Outcome | Not modeled | Observed variables and environment | Trait/structure terms | Trait (TO/PO), direction, value + unit, comparator, conditions |
| Evidence | — | — | — | Location in the publication, optional text span |
| Status | Community standard | Community standard | Reference ontologies | Proposal for community evaluation; schema + validator provided |

#### Supplementary Table S6: Record-level vs article-level shares (paper-level sensitivity)

| **dimension** | **category** | **records** | **% of 14,358 records** | **articles** | **% of 6,452 articles** |
| --- | --- | --- | --- | --- | --- |
| modification class | Overexpression / ectopic expression | 8424 | 58.7 | 4592 | 71.2 |
| modification class | Heterologous / foreign-gene expression | 1578 | 11.0 | 970 | 15.0 |
| modification class | Gene editing (CRISPR / TALEN / ZFN) | 1205 | 8.4 | 769 | 11.9 |
| modification class | Knockout / knockdown | 1121 | 7.8 | 803 | 12.4 |
| modification class | Complementation | 865 | 6.0 | 645 | 10.0 |
| modification class | RNAi / silencing | 778 | 5.4 | 596 | 9.2 |
| modification class | Promoter / reporter / localization | 210 | 1.5 | 138 | 2.1 |
| modification class | Other / unclassified | 154 | 1.1 | 98 | 1.5 |
| modification class | Mutant / point mutation | 23 | 0.2 | 15 | 0.2 |
| trait class | disease | 1365 | 9.5 | 870 | 13.5 |
| trait class | flowering | 1098 | 7.6 | 692 | 10.7 |
| trait class | yield | 1029 | 7.2 | 723 | 11.2 |
| trait class | repro | 887 | 6.2 | 557 | 8.6 |
| trait class | pigment | 870 | 6.1 | 598 | 9.3 |
| trait class | photo | 704 | 4.9 | 511 | 7.9 |
| trait class | salt | 670 | 4.7 | 438 | 6.8 |
| trait class | growth | 632 | 4.4 | 444 | 6.9 |
| trait class | root | 619 | 4.3 | 411 | 6.4 |
| trait class | drought | 578 | 4.0 | 412 | 6.4 |
| trait class | nutrient | 456 | 3.2 | 293 | 4.5 |
| trait class | germination | 423 | 2.9 | 306 | 4.7 |
| trait class | oilseed | 383 | 2.7 | 255 | 4.0 |
| trait class | cold | 316 | 2.2 | 207 | 3.2 |
| trait class | pest | 229 | 1.6 | 159 | 2.5 |
| trait class | heat | 202 | 1.4 | 120 | 1.9 |
| trait class | senescence | 177 | 1.2 | 115 | 1.8 |
| trait class | herbicide | 122 | 0.8 | 89 | 1.4 |
| host species | Arabidopsis thaliana | 6164 | 42.9 | 2647 | 41.0 |
| host species | Oryza sativa | 1935 | 13.5 | 927 | 14.4 |
| host species | Nicotiana benthamiana | 1156 | 8.1 | 541 | 8.4 |
| host species | Nicotiana tabacum | 784 | 5.5 | 472 | 7.3 |
| host species | Solanum lycopersicum | 609 | 4.2 | 344 | 5.3 |
| host species | Triticum aestivum | 281 | 2.0 | 153 | 2.4 |
| host species | Zea mays | 272 | 1.9 | 164 | 2.5 |
| host species | Glycine max | 246 | 1.7 | 143 | 2.2 |
| host species | Populus | 233 | 1.6 | 130 | 2.0 |
| host species | Gossypium hirsutum | 187 | 1.3 | 120 | 1.9 |
| host species | Solanum tuberosum | 152 | 1.1 | 107 | 1.7 |
| host species | Malus domestica | 118 | 0.8 | 61 | 0.9 |
| host species | Hordeum vulgare | 109 | 0.8 | 66 | 1.0 |
| host species | Brassica napus | 87 | 0.6 | 58 | 0.9 |
| host species | Physcomitrium patens | 85 | 0.6 | 37 | 0.6 |
| TF family | WRKY | 268 | 1.9 | 137 | 2.1 |
| TF family | MYB | 188 | 1.3 | 123 | 1.9 |
| TF family | bHLH | 161 | 1.1 | 85 | 1.3 |
| TF family | ERF | 127 | 0.9 | 78 | 1.2 |
| TF family | MIKC_MADS | 116 | 0.8 | 62 | 1.0 |
| TF family | NAC | 112 | 0.8 | 70 | 1.1 |
| TF family | bZIP | 97 | 0.7 | 56 | 0.9 |
| TF family | ARF | 65 | 0.5 | 38 | 0.6 |
| TF family | C2H2 | 59 | 0.4 | 25 | 0.4 |
| TF family | GRAS | 54 | 0.4 | 31 | 0.5 |

**Notes.** Spearman rank correlation between record and article counts: modification class 0.983, trait class 0.99, host species 0.993, TF family 0.976. Gene–trait edges: 7,923; 1,379 have ≥2 records from a single article and 388 are supported by ≥2 articles (record vs article edge weight, Spearman 0.449). Host species are counted as in Figure 1B: every host named in a record counts, so a record or article can count toward more than one host, and Populus is a genus-level aggregate.

#### Supplementary Table S7: Regulatory-part classes in the released construct strings

| **Class** | **Records** |
| --- | --- |
| CaMV35S | 5,268 |
| Ubiquitin | 688 |
| U6/U3 Pol III | 385 |
| native/own | 360 |
| inducible (Dex/XVE/HS) | 129 |
| seed-specific | 67 |
| RbcS/Cab light | 61 |
| Actin | 24 |
| rd29A stress-inducible | 14 |
| root-specific | 1 |
| NOS terminator | 1,074 |
| 35S terminator | 124 |
| OCS terminator | 66 |
| rbcS terminator | 1 |

**Notes.** Root-preferential promoters: the published class matched only the literal words 'root-specific' (1 record). Extending the class with named root-preferential promoters (RCc3, TobRB7/RB7, Pyk10, rolD, SCR, SHR, WOX5, EXPA7, PHT1, PLT, LRP1, SlREO) gives 23 records with such a promoter in a promoter slot. 855 records have both a promoter class and a terminator class. Terminator counts are those after the terminator check (Supplementary Note 3).

#### Supplementary Table S8: Screening-retained vs contributing articles by publication year

| **publication_year** | **sampled_retained_articles** | **estimated_retained_articles(all)** | **release_contributing_articles** | **contributing/retained** |
| --- | --- | --- | --- | --- |
| 2015 | 147 | 3185 | 394 | 12.4% |
| 2016 | 183 | 3965 | 466 | 11.8% |
| 2017 | 173 | 3748 | 453 | 12.1% |
| 2018 | 177 | 3835 | 400 | 10.4% |
| 2019 | 194 | 4203 | 488 | 11.6% |
| 2020 | 211 | 4572 | 371 | 8.1% |
| 2021 | 247 | 5352 | 133 | 2.5% |
| 2022 | 209 | 4528 | 109 | 2.4% |
| 2023 | 234 | 5070 | 432 | 8.5% |
| 2024 | 306 | 6630 | 671 | 10.1% |
| 2025 | 343 | 7432 | 737 | 9.9% |
| 2026 | 127 | 2752 | 234 | 8.5% |

**Notes.** Retained articles were estimated from a seeded random sample of 3,128 retained accessions with a JATS publication year. Screening retained as many 2021–2022 articles as adjacent years (2.5% and 2.4% of retained articles contributed records, versus 8–12% in other years).

#### Supplementary Table S9.1: Time-split test of graph-based hypothesis generation (cutoff 2022; test records 2023–2026)

| **Task** | **Score** | **AUROC (95% CI)** | **AUPRC (95% CI)** | **AUPRC / base rate** | **P@20** | **P@50** | **Hits in top 100** | **R@200** |
| --- | --- | --- | --- | --- | --- | --- | --- | --- |
| Cross-species transfer | Graph rule (pre-specified) | 0.754 (0.698–0.815) | 0.041 (0.018–0.063) | 3.5 | 0.20 | 0.14 | 7 | 0.07 |
| Cross-species transfer | Graph model | 0.904 (0.867–0.940) | 0.138 (0.094–0.228) | 11.7 | 0.10 | 0.20 | 20 | 0.23 |
| Cross-species transfer | Research-intensity baseline | 0.898 (0.858–0.932) | 0.139 (0.087–0.208) | 11.8 | 0.20 | 0.28 | 23 | 0.23 |
| Cross-species transfer | Research-intensity model | 0.901 (0.862–0.936) | 0.142 (0.093–0.227) | 12.1 | 0.25 | 0.20 | 26 | 0.23 |
| Cross-species transfer | Random | 0.497 (0.457–0.537) | 0.013 (0.007–0.019) | 1.1 | 0.05 | 0.04 | 3 | 0.02 |
| Gene–trait association | Graph rule (pre-specified) | 0.646 (0.592–0.693) | 0.031 (0.021–0.045) | 2.1 | 0.05 | 0.08 | 8 | 0.04 |
| Gene–trait association | Graph model | 0.778 (0.735–0.815) | 0.066 (0.047–0.089) | 4.4 | 0.05 | 0.12 | 11 | 0.08 |
| Gene–trait association | Research-intensity baseline | 0.766 (0.729–0.801) | 0.061 (0.043–0.085) | 4.1 | 0.05 | 0.14 | 8 | 0.07 |
| Gene–trait association | Research-intensity model | 0.782 (0.740–0.819) | 0.062 (0.044–0.085) | 4.2 | 0.05 | 0.10 | 10 | 0.07 |
| Gene–trait association | Random | 0.515 (0.484–0.545) | 0.016 (0.013–0.020) | 1.1 | 0.00 | 0.02 | 1 | 0.02 |

**Notes.** Candidates and positives: cross-species transfer 13,793 and 162 (base rate 0.012); gene–trait association 17,696 and 261 (0.015). Of the links first reported after 2022, 162 of 378 transfers and 261 of 501 gene–trait links were candidates at the cutoff; the others involved gene families, orthogroups, hosts or trait combinations not yet in the graph. AUROC, area under the ROC curve; AUPRC, average precision; AUPRC / base rate, AUPRC divided by the share of candidates that were positive; P@k, share of the k top-ranked candidates reported later; R@200, share of all positives among the 200 top-ranked candidates.

#### Supplementary Table S9.2: Paired differences between scores in the time-split test (cutoff 2022; test records 2023–2026)

| **Task** | **Comparison** | **Metric** | **Difference** | **95% CI** |
| --- | --- | --- | --- | --- |
| Cross-species transfer | Graph model − Research-intensity model | AUROC | 0.003 | -0.002 to 0.011 |
| Cross-species transfer | Graph model − Research-intensity model | AUPRC | -0.004 | -0.019 to 0.015 |
| Cross-species transfer | Graph rule (pre-specified) − Research-intensity baseline | AUROC | -0.145 | -0.193 to -0.086 |
| Cross-species transfer | Graph rule (pre-specified) − Research-intensity baseline | AUPRC | -0.098 | -0.176 to -0.044 |
| Cross-species transfer | Graph model − Random | AUROC | 0.407 | 0.360 to 0.453 |
| Cross-species transfer | Graph model − Random | AUPRC | 0.125 | 0.082 to 0.218 |
| Cross-species transfer | Graph rule (pre-specified) − Random | AUROC | 0.257 | 0.210 to 0.315 |
| Cross-species transfer | Graph rule (pre-specified) − Random | AUPRC | 0.028 | 0.008 to 0.049 |
| Gene–trait association | Graph model − Research-intensity model | AUROC | -0.004 | -0.016 to 0.008 |
| Gene–trait association | Graph model − Research-intensity model | AUPRC | 0.003 | -0.004 to 0.012 |
| Gene–trait association | Graph rule (pre-specified) − Research-intensity baseline | AUROC | -0.120 | -0.163 to -0.077 |
| Gene–trait association | Graph rule (pre-specified) − Research-intensity baseline | AUPRC | -0.029 | -0.051 to -0.010 |
| Gene–trait association | Graph model − Random | AUROC | 0.262 | 0.209 to 0.309 |
| Gene–trait association | Graph model − Random | AUPRC | 0.050 | 0.033 to 0.071 |
| Gene–trait association | Graph rule (pre-specified) − Random | AUROC | 0.131 | 0.066 to 0.193 |
| Gene–trait association | Graph rule (pre-specified) − Random | AUPRC | 0.015 | 0.006 to 0.029 |

**Notes.** Paired differences use the same 1,000 bootstrap resamples. AUROC and AUPRC are defined in the notes to Supplementary Table S9.1. The sensitivity analysis (positives confirmed by at least two articles) gave the same conclusions; all values are in Supplementary Data 3 (sheets metrics and paired_differences).

### Supplementary Figures

#### Supplementary Figure S1: Audited processing flow

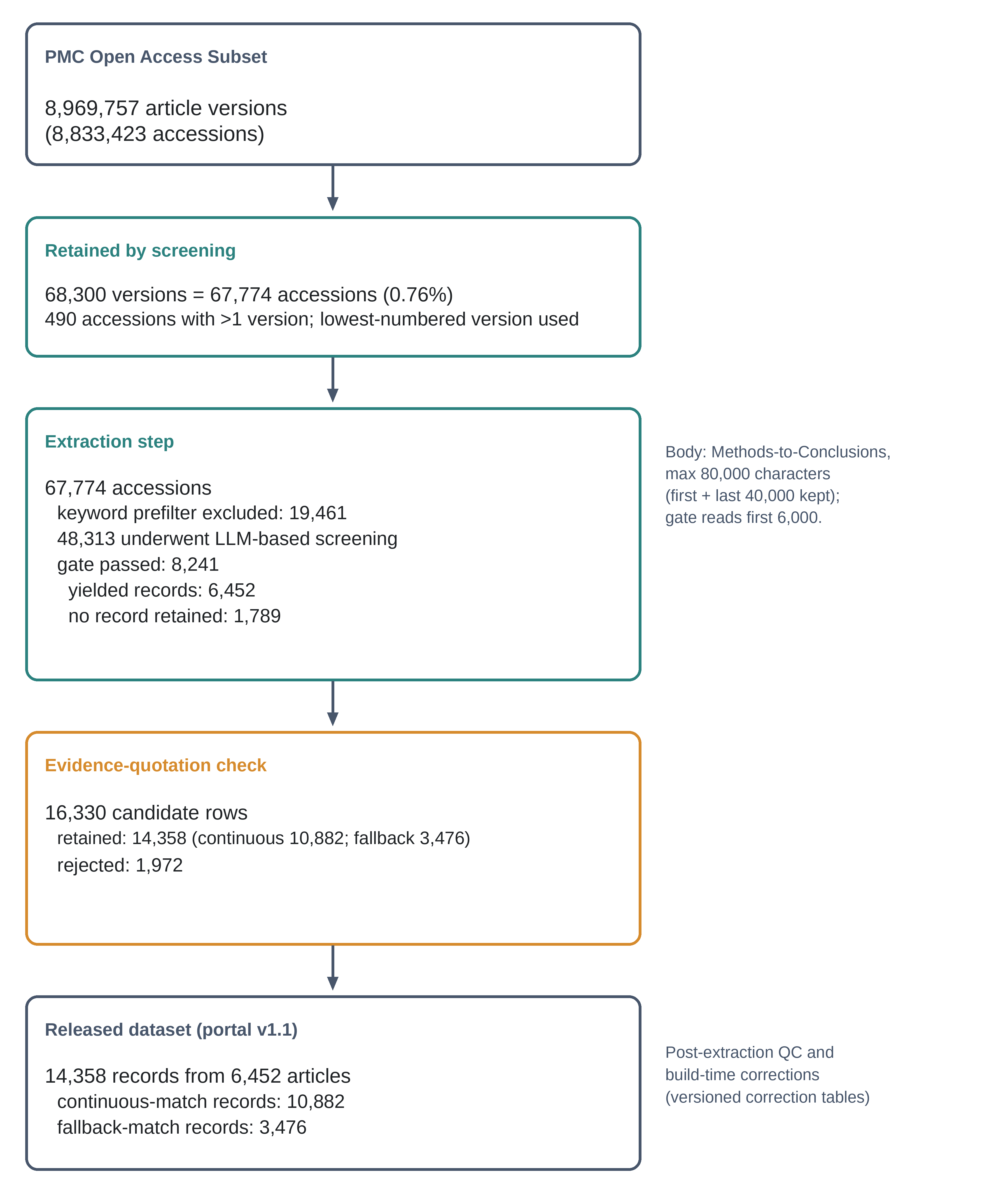

### Supplementary Data

Supplementary Data 1–3 are machine-readable files that cannot be reproduced in this document. They are deposited in the public repository https://github.com/khanaliyawar4/Plant_Atlas.

| **Item** | **Content** | **Location in the repository** |
| --- | --- | --- |
| Supplementary Data 1 | Reporting-profile JSON Schema (v0.1.0), a valid and an invalid worked example, and the validator; reproduced in Supplementary Note 6 | reporting_profile/ |
| Supplementary Data 2 | Source code for screening, extraction, annotation, portal build and analysis, with prompts, schemas and lexicons | The repository |
| Supplementary Data 3 | Time-split test: metrics, paired differences and the 50 highest-ranked untested candidates for each task (.xlsx) | supplementary_data/Supplementary_Data_3_time_split_test.xlsx |
